# PrimateFace: A Machine Learning Resource for Automated Face Analysis in Human and Non-human Primates

**DOI:** 10.1101/2025.08.12.669927

**Authors:** Felipe Parodi, Jordan K. Matelsky, Alessandro P. Lamacchia, Melanie Segado, Yaoguang Jiang, Alejandra Regla-Vargas, Liala Sofi, Clare Kimock, Bridget M. Waller, Michael L. Platt, Konrad P. Kording

## Abstract

Machine learning has revolutionized human face analysis, but equivalent tools for non-human primates remain limited and species-specific, hindering progress in neuroscience, anthropology, and conservation. Here, we present PrimateFace, a comprehensive, cross-species platform for primate facial analysis comprising a systematically curated dataset of 260,000+ images spanning over 60 genera, including a genus-balanced subset of 60,000 images, annotated with bounding boxes and facial landmark configurations. Face detection and facial landmark estimation models trained on PrimateFace achieve high cross-species performance, from tarsiers to gorillas, achieving performance comparable to baseline models trained exclusively on human data (0.34 vs. 0.39 mAP for face detection; 0.061 vs. 0.053 normalized landmark error), demonstrating the generalization benefits of cross-species training. PrimateFace enables diverse downstream applications including individual recognition, gaze analysis, and automated extraction of stereotyped (e.g., lip-smacking) and subtle (e.g., soft left turn) facial movements. PrimateFace provides a standardized platform for facial phenotyping across the primate order, empowering data-driven studies that advance the health and well-being of human and non-human primates. All models, notebooks, and data can be found at github.com/KordingLab/PrimateFace.

---

Faces serve as essential conduits for conveying information critical to social animals, including identity, emotions, intentions, focus, and social status^1,2^. This is particularly true for human and non-human primates^3^, with relevance to psychology, neuroscience, evolutionary biology, anthropology, and conservation^2,4,5^. Despite its profound scientific and practical importance, our ability to quantitatively decode communication in non-human primate (NHP) faces remains surprisingly limited^6^.

A primary obstacle lies in the remarkable diversity of non-human primate facial morphology, fur coloration, and facial muscle size, kinematics, and motor control^4,5^. This variation, coupled with complex natural environments, makes manual analysis of facial movements exceedingly time-consuming and prone to subjective biases. While systems like the Facial Action Coding System (FACS) and its primate-specific adaptations (e.g., ChimpFACS, MaqFACS, CalliFACS) provide valuable frameworks for objective measurement^3^, manual coding of these action units remains labor-intensive, hindering progress across primate behavioral research^4,5,7^. This limitation highlights the critical need for foundational computational infrastructure – robust face detection and facial landmark (also known as face ‘pose’ or ‘keypoints’) estimation – that could enable automated analysis at the scale and precision necessary to advance understanding of primate facial communication.

Recent advances in deep learning have revolutionized human facial analysis, achieving remarkable precision in face detection, landmark tracking, and emotion recognition^8^. Datasets like WIDERFace and COCO-WholeBody have driven this progress by providing vast quantities of labeled data to train sophisticated algorithms^9,10^. The remarkable success of these methods for human facial analysis raises the tantalizing prospect of leveraging similar approaches to automate the analysis of non-human primate faces. Despite the clear parallels and potential benefits, a significant technological gap persists between human and non-human primate facial analysis systems.

Current computational primatology efforts prioritize individual recognition^11^ or full-body pose estimation^12–16^, while existing facial analysis approaches are typically species-specific^17–25^, trained on small datasets, and lack the taxonomic breadth required for robust cross-species analysis^26–30^ (**Extended Data Table 1**). Despite serving as the foundation for downstream applications, including cross-species comparisons^31^ and pain assessment^32^, no practical cross-primate solution exists. This disparity stems from three fundamental challenges: 1) immense annotation costs associated with diverse NHP faces; 2) significant morphological divergence causing poor generalization when applying human-specific models to NHP data; and 3) scarcity of large-scale, publicly available cross-species face data.

To address this gap, we developed **PrimateFace.** Unlike existing species-specific solutions, PrimateFace provides immediate cross-species capability, integrating: (1) a large-scale and taxonomically diverse dataset of over 260,000 images across 60+ genera, annotated with face bounding boxes and dual (48-keypoint NHP-centric and 68-keypoint standard) facial landmark configurations; (2) a suite of deep learning models that achieve high-performance, cross-species face detection and facial landmark estimation; and (3) an open-source ecosystem of software and tutorials to facilitate scientific applications. While trained on static images, PrimateFace models apply frame-by-frame to video data, enabling temporal analysis of facial behaviors.

We demonstrate that PrimateFace models generalize effectively from tarsiers to gorillas – and even to human face benchmarks – enabling applications including individual recognition, gaze analysis, and automated discovery of stereotyped (e.g., lip-smacking) facial movements. PrimateFace provides a standardized platform for facial phenotyping across the primate order, empowering data-driven studies that advance both human and non-human behavior, cognition, and welfare.

## Results

### PrimateFace is an integrated ecosystem for primate face analysis

PrimateFace addresses the need for dependable, cross-species primate facial analysis by providing a suite of resources (**Fig. 1**), including a scalable data curation and annotation workflow (**Fig. 1**). To develop PrimateFace, we first curated several publicly available primate datasets (**Extended Data Table 1**). Following curation, we employed DINOv2-guided image sampling (**Di**stillation of k**no**wledge^33^) to ensure diverse representation from large-scale, unlabeled image pools for rapid, semi-automated annotation in our custom graphical user interface (GUI). This GUI facilitates rapid, iterative labeling by allowing human annotators to efficiently review and refine pseudo-labels of face boxes or landmarks generated by progressively improving PrimateFace models (**Fig. 1c, Extended Data Fig. 3c**). The ecosystem additionally incorporates vision-language models (e.g., Gemini 2.5 Pro) for semi-automated genus classification with human-in-the-loop correction, providing valuable taxonomic organization capabilities for large-scale datasets. PrimateFace is developed as a growing ecosystem and we provide a community contribution portal to foster collaborative expansion.

**Figure 1.**
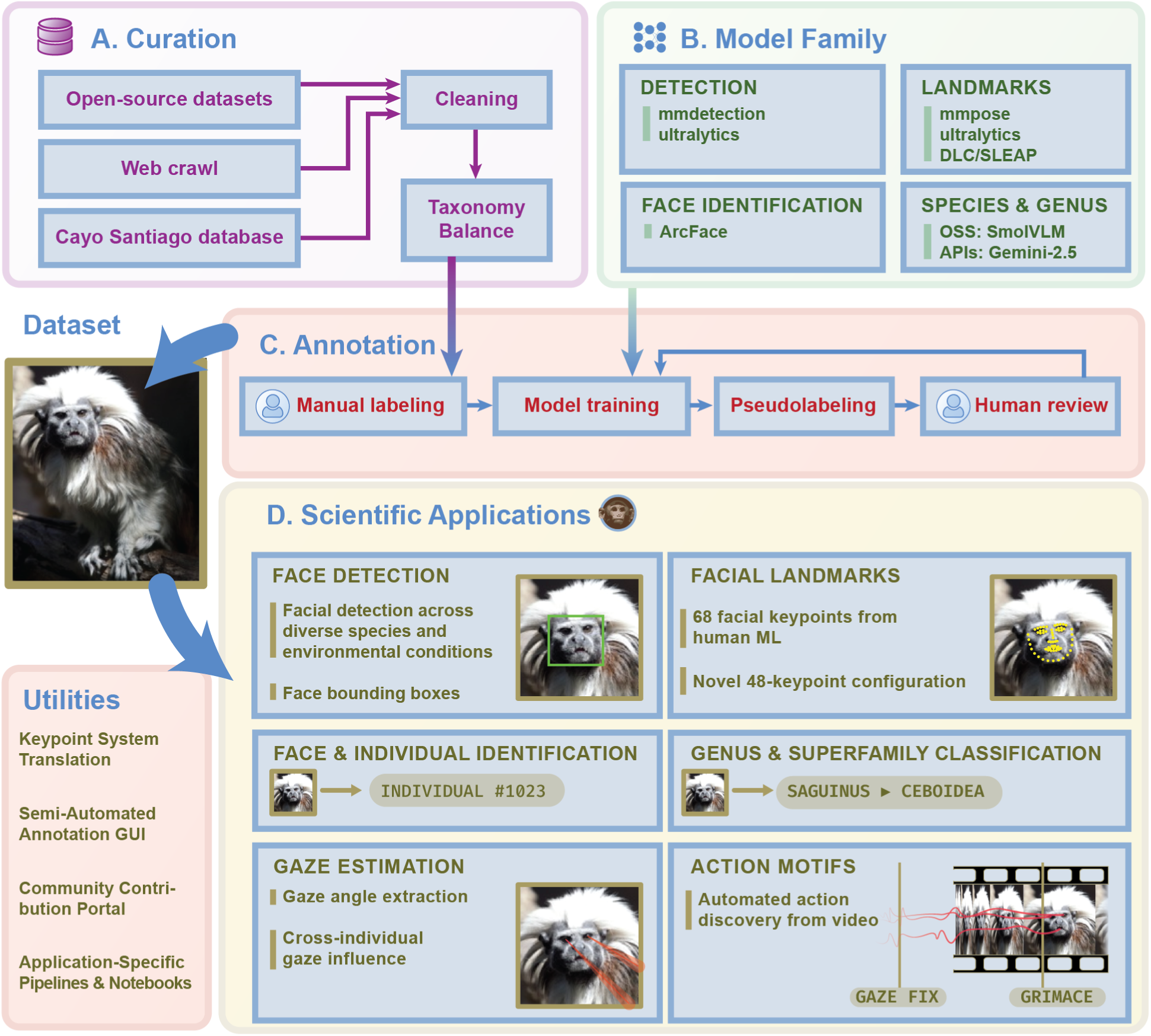
PrimateFace is an integrated ecosystem for primate facial analysis. PrimateFace unifies dataset curation, model development, annotation, and inference. **A.** Primate facial images were aggregated from open-source datasets, targeted web scraping, and institutional repositories (e.g., Cayo Santiago Field Station), followed by cleaning and taxonomy-balancing. **B.** Detection, landmark estimation, identity, and species/genus classification models were developed across multiple open-source frameworks (MMDetection, MMPose, Ultralytics, DeepLabCut, SLEAP, InsightFace, and HuggingFace). **C.** Data labeling was accelerated via a hybrid workflow combining manual annotation, iterative pseudo-labeling, and human-in-the-loop review. **D.** The trained models support core tasks including face detection, facial landmark localization, individual recognition, facial movement extraction, and genus classification – enabling robust, automated primate face analysis. Together, these stages yield a scalable, annotation-efficient workflow and a unified resource for primate face analysis.

This structured approach to data curation has yielded the PrimateFace dataset, the primary deliverable. The dataset currently encompasses over 260,000 images spanning 60+ primate genera, annotated with bounding boxes and both 48-point (NHP-centric) and 68-point (human-standard) facial landmarks (**Fig. 2**), exportable to common formats like SLEAP^15^ and DeepLabCut^16^ (**Suppl Fig. 2**).

**Figure 2.**
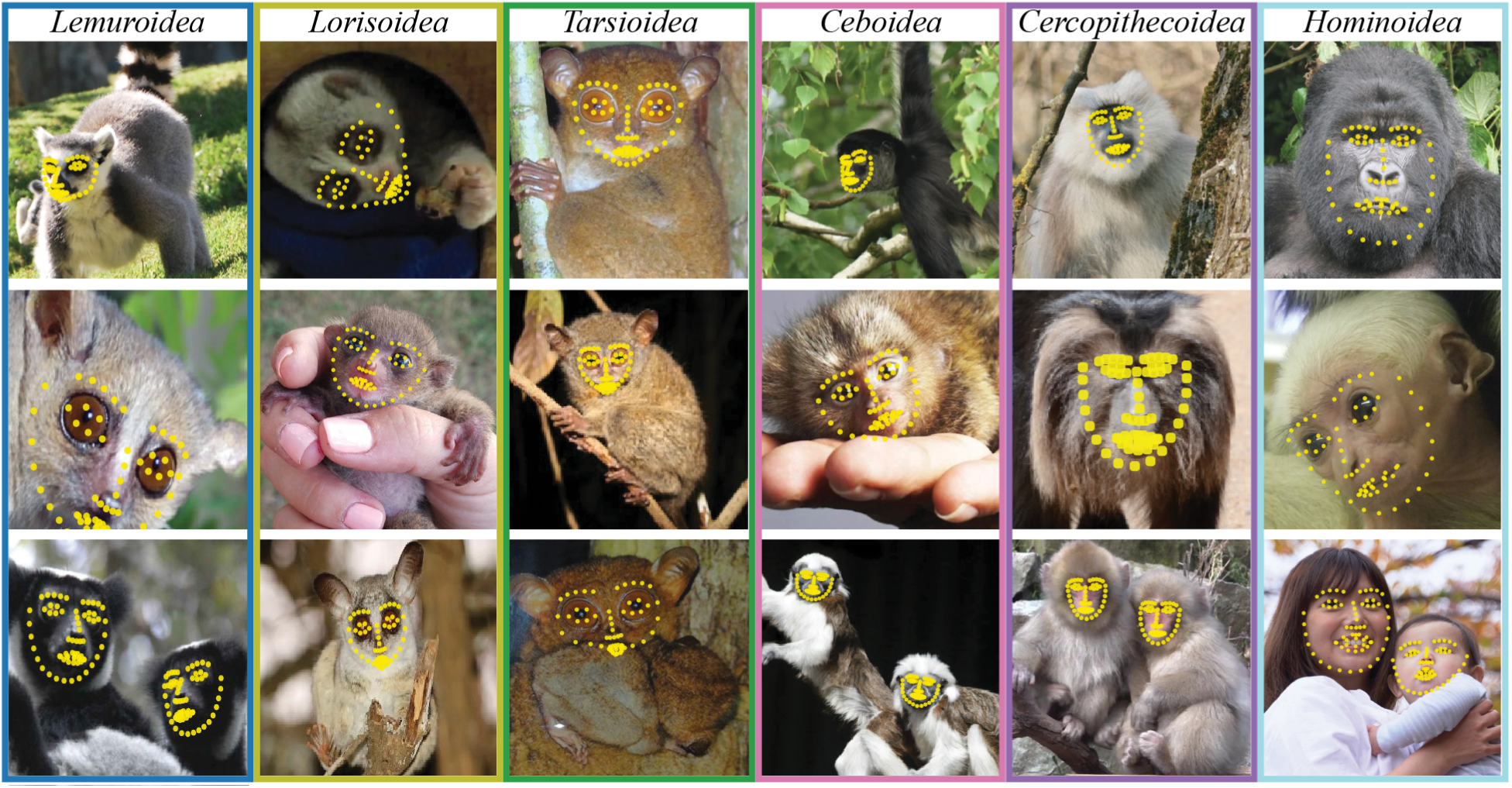
PrimateFace dataset provides a diverse foundation for cross-species analysis. Representative annotated samples from the six primate superfamilies: Lemuroidea, Lorisoidea, Tarsioidea, Ceboidea, Cercopithecoidea, and Hominoidea. Each image is labeled with 68 facial landmarks. The dataset spans diverse species, developmental stages (infant to adult), environments (lab, zoo, wild), and social contexts (solitary, dyadic, group-living), capturing both common and rare phenotypes under naturalistic lighting conditions. PrimateFace provides a foundation for cross-species benchmarking and the development of generalizable primate face analysis models. Images depicting primate handling represent standard veterinary or husbandry care in zoological settings. **Top row genera:** Lemur, Pseudopotto, Tarsius, Ateles, Semnopithecus, Gorilla. **Middle row:** Mirza, Galago, Cephalopachus, Cebuella, Macaca, Hylobates. **Bottom row:** Lemur, Paragalago, Carlito, Callithrix, Macaca, Homo.

Built upon this dataset, PrimateFace delivers a suite of pre-trained models for high-performance, cross-species face detection (**Fig. 3**) and facial landmark estimation (**Fig. 5**). Remarkably, face detection models trained solely on PrimateFace achieve 0.34 mAP@0.50 (standard detection accuracy metric where higher values indicate better performance) on the challenging WIDERFace human benchmark^9^ – within 0.05 of human-face baselines (0.39 mAP@0.50) (**Fig. 3c**).

**Figure 3.**
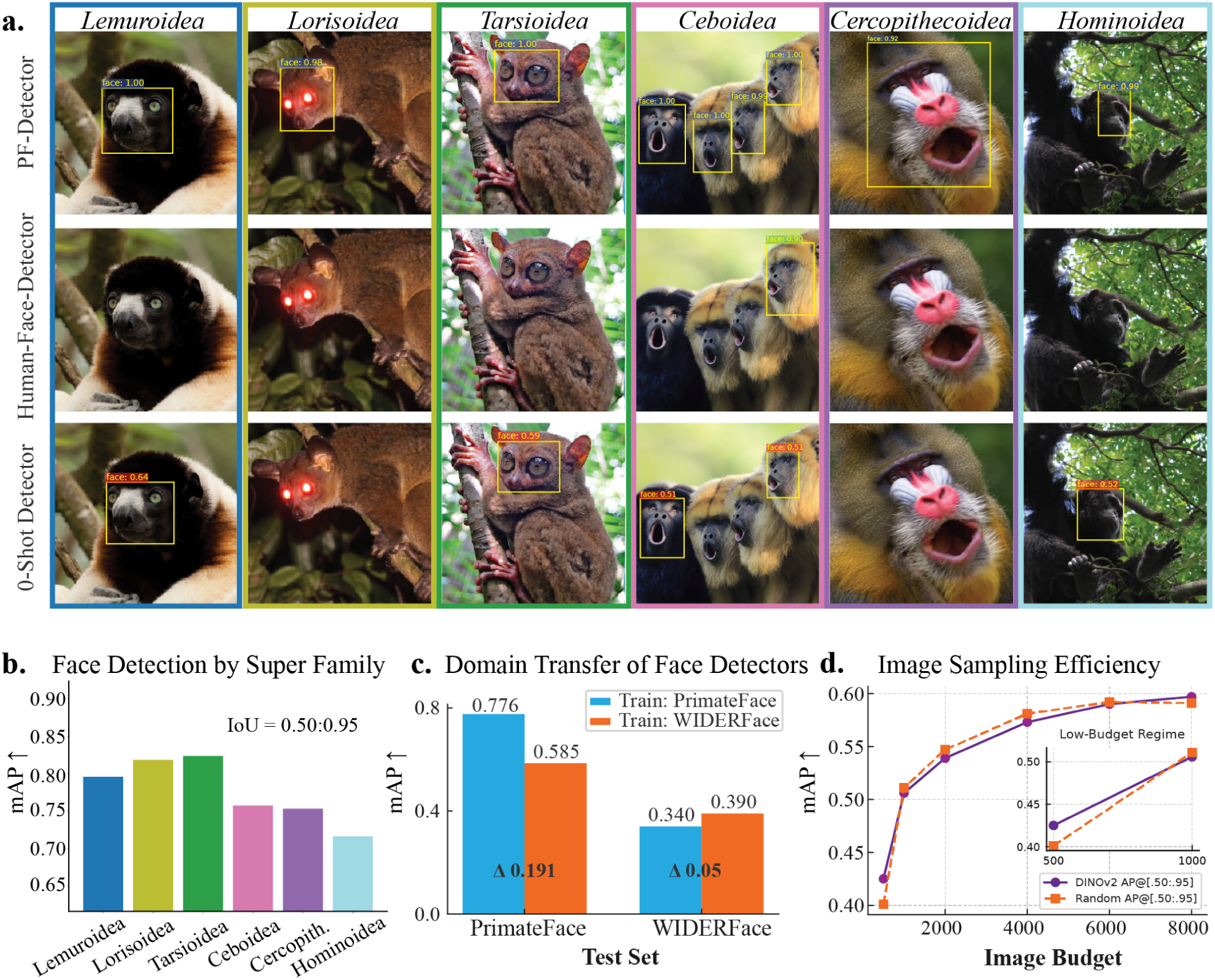
Evaluation of PrimateFace detection models. **A.** Qualitative comparison of a PrimateFace-finetuned face detection model (top); an off-the-shelf human face detector (middle); and an ‘open-vocabulary’, zero-shot detection model prompted with ‘Face’. PrimateFace and human-face models were thresholded at 0.75 confidence; the zero-shot model was thresholded at 0.5. **B.** Face detection performance by superfamily. Models trained on genus-balanced PrimateFace data exhibit highest performance on Tarsioidea and lowest on Hominoidea. **C.** PrimateFace-trained detectors achieve high accuracy on both PrimateFace and WIDERFace test sets, demonstrating effective transfer between primate and human face detection tasks. **D.** DINOv2-guided sampling shows improved data efficiency over random sampling in the low-data regime (inset: 500, 1000 image performance), with peak advantage around 500 training images. Performance converges at larger dataset sizes, demonstrating the utility of guided sampling specifically for annotation-constrained scenarios typical when working with rare primate taxa. **Acronyms**: PF, PrimateFace (dataset); mAP, mean average precision; Cercopith., Cercopithecoidea superfamily.

This cross-species advantage extends to facial landmark estimation, where PrimateFace models achieve 0.061 normalized mean error (NME) (a size-independent accuracy measure where lower values indicate more precise landmark prediction), on COCO-WholeBody-Face^10^, differing by only 0.008 from human-face-trained models (0.053 NME) (**Fig. 5c**). Critically, this generalization is unidirectional – human-face models show substantial performance degradation when applied to primate faces, underscoring the value of morphologically diverse training data for cross-species applications. Finally, to enhance interoperability between common Python libraries, the ecosystem includes a dedicated workflow for converting between the 68- and 48-landmark configurations (**Fig. 4**).

**Figure 4.**
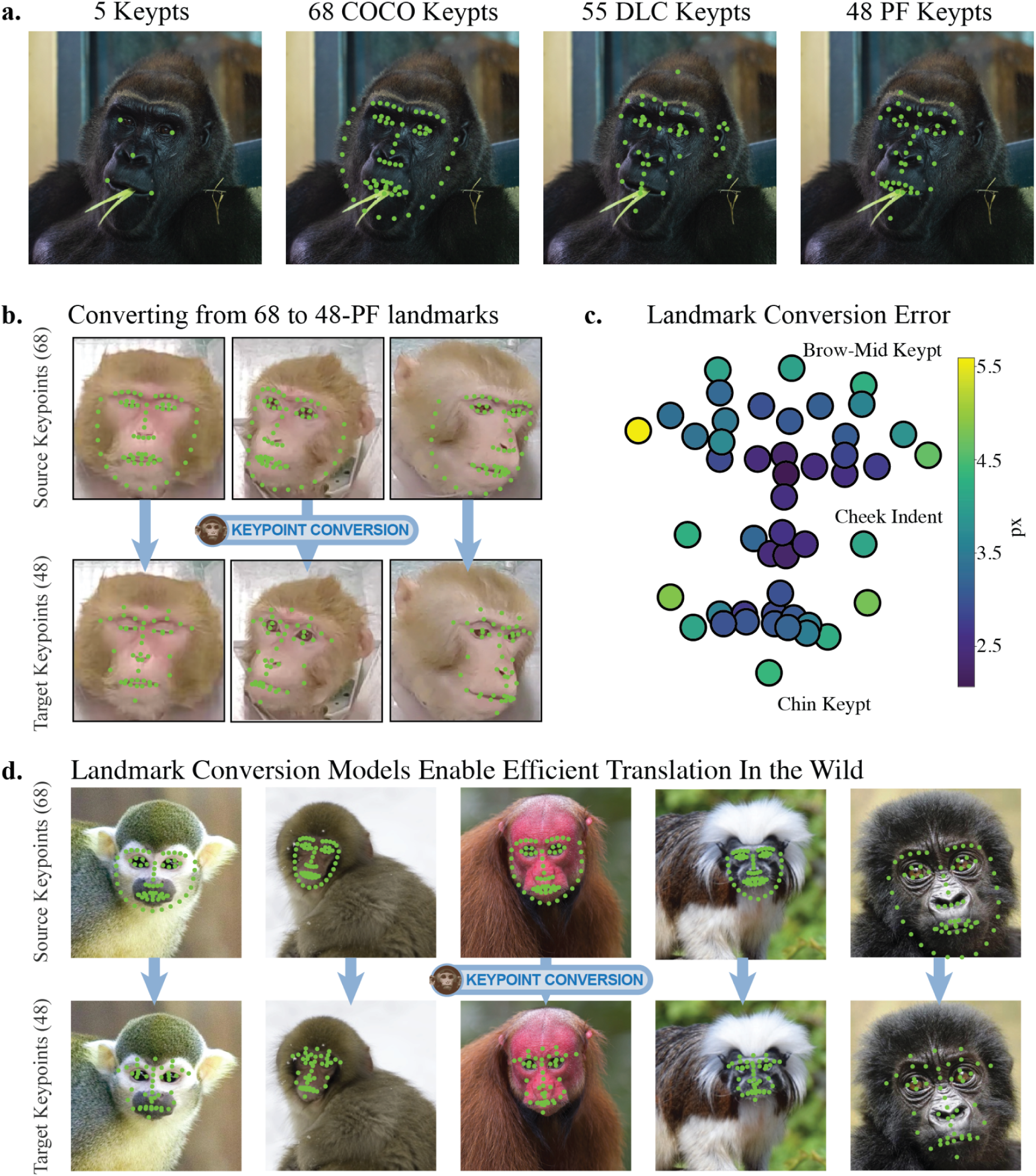
PrimateFace enables landmark conversion across facial landmark conventions. **A.** Common face landmark conventions include 5-landmarks (far left); COCO 68-landmarks (second from the left); DeepLabCut MacaqueFace 55-landmarks (second from the right); and the PrimateFace custom 48-landmark model (far right), which is a subset of the COCO-68 and DLC-55 models. **B.** Training scheme for the facial landmark conversion model, which enables translation between landmark schemes. Models were trained on laboratory-collected high-resolution (4K) images of macaques. **C.** Pixel error heatmap for the 48-predicted landmarks in 4K-resolution images. Darker color indicates lower error. **D.** Example landmark conversions on held-out images. **Acronyms**: Kpt, keypoint (referring to the facial landmarks).

**Figure 5.**
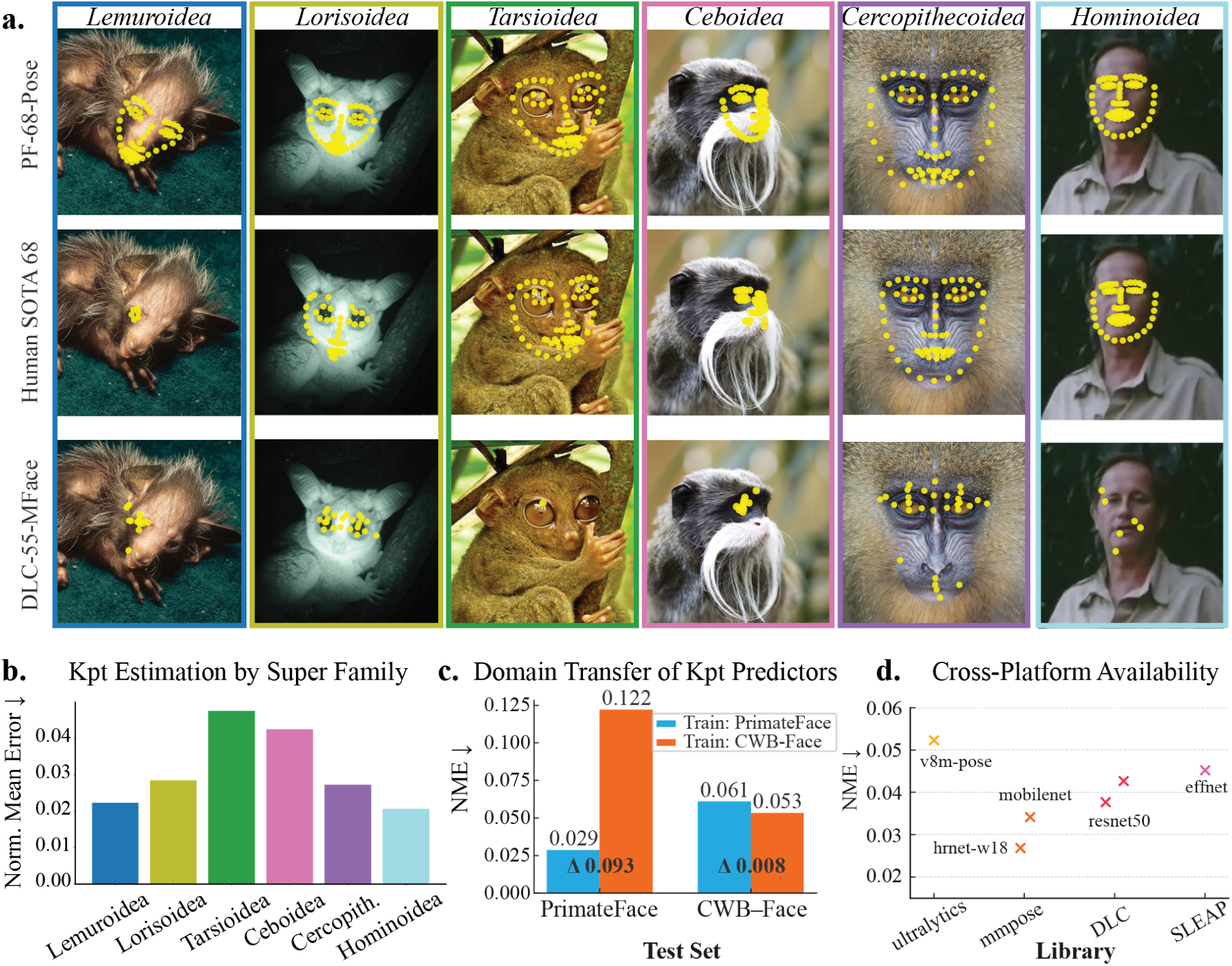
PrimateFace pretrained models enable species-agnostic landmark estimation. **A.** Qualitative comparison of a PrimateFace-finetuned facial landmark estimation model (top); an off-the-shelf human face detection model (middle); and the pretrained DeepLabCut MacaqueFace (55-keypoint) model available in Model Zoo, originally trained on macaque images^16,22^. All models used the same PF-trained Cascade-RCNN face detector. Models were thresholded at 0.75 confidence. **B.** Facial landmark estimation performance by superfamily. Models trained on genus-balanced PrimateFace data exhibit lowest performance for genera for which we have little data (e.g., Tarsiers) and highest on images containing Hominoidea and Lemuroidea. **C.** Training on PrimateFace achieves competitive performance on the challenging human face dataset COCO-WholeBody-Face^10^. **D.** Pre-trained PF-models are available in DeepLabCut, SLEAP, Ultralytics, or mmpose for smooth integration into existing research. **Acronyms**: Kpt, keypoint (or landmarks); SOTA, state-of-the-art; CWB, COCO-WholeBody-Face; DLC, DeepLabCut; MFace, Macaque Face; Norm., normalized; effnet, EfficientNet.

PrimateFace enables robust quantification of dynamic facial behaviors by tracking facial landmarks over time in video sequences. The system outputs precise coordinate positions of facial landmarks – such as mouth corners, eye centers, and nose tip – across video frames, enabling researchers to calculate movement velocities, distances between features, and temporal patterns of facial behavior. This provides objective, quantitative measurements for behaviors that previously relied on subjective visual assessment. However, PrimateFace does not automatically classify discrete facial expressions or emotions – it provides the precise movement data that researchers can then analyze and code according to their specific research questions. The five *Scientific Applications* demonstrate this capability in action, showing how landmark tracking enables automated individual recognition, vocal-motor coupling analysis, social attention dynamics, and discovery of subtle behavioral patterns across primates.

### PrimateFace dataset provides a foundation for cross-species face analysis

Prior to PrimateFace, no dataset captured the vast morphological and expressive diversity of primates. The PrimateFace dataset encompasses over 260,000 images: an extensive aggregation of 200,000+ images sourced and re-annotated from diverse public datasets (see **Extended Data Table 1** for full list of sources), augmented by a novel collection of 60,000 images sampled for taxonomic balance across 60+ primate genera. This effort uniquely extends coverage across the primate order, including previously underrepresented groups such as tarsiers, alongside robust representation of the superfamilies: Lemuroidea, Lorisoidea, Tarsioidea, Ceboidea, Cercopithecoidea, and Hominoidea^34^. PrimateFace captures this taxonomic breadth across a wide array of ecologically-relevant conditions, including images from wild, captive, and laboratory environments, depicting a spectrum of naturalistic social interactions, developmental stages from infancy to adulthood, and diverse facial expressions, all under varied lighting and image quality (**Fig. 2, Suppl Fig. 1**). This combination of scale, taxonomic balancing, and contextual richness establishes PrimateFace as a uniquely versatile dataset, poised to unlock frontiers in the automated quantitative study of primate facial behavior.

Achieving comprehensive taxonomic coverage across 60+ genera presented a critical sampling challenge. Traditional random sampling approaches are prohibitively inefficient for taxonomically imbalanced datasets, often requiring full annotation before capturing sufficient diversity within rare taxa like tarsiers or aye-ayes. To address this, we leveraged DINOv2^33^, a self-supervised vision transformer that captures semantic visual features, to identify and prioritize representationally diverse images across the primate order (**Extended Data Fig. 3**). Images grouped by DINOv2 image feature embeddings frequently corresponded to distinct genera or species, validating that the approach captured meaningful visual cross-primate diversity (**Extended Data Fig. 3b, Extended Data Fig. 4**). This allowed strategic sampling that maximized taxonomic and morphological diversity with minimal annotation overhead.

Empirical validation demonstrated advantages over random sampling in annotation-constrained scenarios. Across multiple labeling budgets (500–8,000 images), Cascade-RCNN face detectors trained on DINOv2-sampled subsets achieved 2-4 point improvements in mAP@[.50:.95] over random sampling, with peak advantage around 500 training images (**Fig. 3d**). This finding informed our dataset construction strategy: we targeted ∼500-850 images per genus to maximize model performance while maintaining taxonomic breadth (**Extended Data Fig. 1**). While both approaches converged at larger dataset sizes, DINOv2 sampling provided essential efficiency gains required for taxonomically diverse, resource-constrained primate research.

This data-driven approach, combined with semi-automated annotation workflows, enabled rapid labeling of face bounding boxes and facial landmarks. Every primate face received a tight bounding box annotation around the face (typically excluding ears), accompanied by both our novel 48-facial-keypoint scheme optimized for cross-species analysis and the standard 68-keypoint format for interoperability with existing pipelines. However, designing landmark schemes that capture morphological diversity from prosimian rostra to great ape facial profiles required careful consideration of existing annotation conventions.

Facial landmark conventions have evolved from rudimentary alignment schemes to increasingly sophisticated anatomical mappings. Early computer vision approaches relied on sparse 5-keypoint configurations sufficient for basic face alignment tasks^8^, while human facial analysis matured around the 68-keypoint standard popularized by datasets like COCO-WholeBody^10^ (**Fig. 4a**, left to right). Recognizing the unique challenges of non-human primate morphology, prior work has developed NHP-specific landmark schemes, most notably DeepLabCut’s 55-keypoint MacaqueFace model in their Model Zoo^16,22^ (**Fig. 4a**, second from the right) – a pioneering effort that enabled essential pseudo-labeling for our annotation pipeline. However, no existing convention adequately captures the morphological heterogeneity spanning the entire primate order, from pronounced baboon muzzles to flat-nosed capuchins.

PrimateFace addresses this gap through rich, dual-layered annotations designed to support both specialized primatological inquiry and broader computer vision applications. The 48-keypoint configuration (**Fig. 4a far right**), derived by combining the most robust landmarks from both DeepLabCut’s MacaqueFace model and the human 68-keypoint standard, strategically emphasizes anatomical landmarks that meet three key criteria: 1) consistent visibility across species (excluding neck nape landmarks often occluded by posture); 2) anatomical precision (omitting COCO-68 chin contours that lack clear anatomical correspondence even in human faces); and 3) universal applicability, namely the exclusion of discrete pupil landmarks since many primate species lack clearly visible pupils. This configuration retains valuable detailed landmarks from the COCO-68 standard, particularly the precise mouth contour and nose bridge annotations essential for facial expression analysis, while ensuring reliable cross-species applicability. Researchers can leverage subsets of these landmarks for region-specific analyses, such as focusing on periocular keypoints for eye movement studies or mouth contours for vocal expression analysis (see *Scientific Application 3*). The parallel 68-keypoint annotations maintain interoperability with existing human facial analysis pipelines, creating a unified foundation for developing future cross-species models for primate behavior analysis. All annotations are exportable to common movement analysis frameworks such as SLEAP^15^, DeepLabCut^16^, OpenMMLab, or Ultralytics. Researchers are encouraged to contribute data at our community contribution portal (primateface.studio).

### PrimateFace pretrained models enable high-performance, cross-species face detection

Accurate face localization across primates represents a fundamental challenge that existing approaches have yet to solve comprehensively. While existing methods achieve species-specific detection, none have demonstrated reliable performance spanning the full taxonomic breadth. PrimateFace-trained detectors address this gap directly, achieving strong face detection across all six primate superfamilies: Lemuroidea, Lorisoidea, Tarsioidea, Ceboidea, Cercopithecoidea, and Hominoidea (**Fig. 3a,b**). We trained multiple state-of-the-art object detection architectures on PrimateFace face bounding box data using default hyperparameters. Models are made available for immediate use through frameworks including MMDetection and Ultralytics.

#### PrimateFace detectors outperform human-face and open-vocabulary models on primate faces

To evaluate the face detectors, we first qualitatively compared PrimateFace-trained detectors against two alternative strategies: 1) human face detectors applied directly to primate images, and 2) *open-vocabulary* detection models that can be prompted with arbitrary object categories. Open-vocabulary (or zero-shot) models like GroundingDINO^35^ represent an emerging paradigm in computer vision, using vision-language pretraining to detect objects specified through natural language prompts (e.g., ‘spider monkey face’) without requiring specialized training. Despite their impressive performance across many domains, these zero-shot models have yet to solve primate face detection, likely due to insufficient representation of in-the-wild primate imagery.

Qualitative comparison across these approaches reveals the advantages of our broad cross-species training approach (**Fig. 3a**). While PrimateFace-trained models consistently detect faces across diverse primate species, human face detectors fail on many non-human primate images, and the open-vocabulary GroundingDINO detector shows inconsistent performance when prompted with “face.” Despite the computer vision field’s increasing emphasis on foundation models capable of zero-shot inference, comprehensive training on morphologically diverse primate facial data remains essential for reliable interspecific detection.

#### Face detection maintains high performance across all primate superfamilies

Beyond qualitative assessment, quantitative evaluation across primate superfamilies confirmed PrimateFace’s species-agnostic performance. Models achieved highest accuracy on Tarsioidea and lowest on Hominoidea (**Fig. 3b**), though all genera maintained detection performance (mAP @0.50:0.95) above 0.60, indicating reliable detection across taxa (**Extended Data Fig. 2a**). Notably, even this lowest performance remained stable, as demonstrated by our models’ competitive performance on human face benchmarks (see cross-domain evaluation **Fig. 3c**). This variation likely reflects differences in data curation and image contexts rather than inherent face detection difficulty: images of Tarsier monkeys typically feature subjects prominently centered with clear facial visibility, whereas Cercopithecoidea and Hominoidea data encompass far greater diversity – from controlled laboratory settings to challenging naturalistic environments with variable lighting, poses, and occlusions. Similar heterogeneity emerged across genera (**Extended Data Fig. 2a**), highlighting how dataset composition and ecological context influence model performance – insights that will inform future cross-species data collection.

#### PrimateFace models generalize effectively to human face detection

Cross-domain evaluation on WIDERFace^9^ revealed that training on diverse primate faces creates generalizable detectors. PrimateFace-trained models achieved 0.340 mAP@0.50 on the challenging WIDERFace human benchmark – within 0.05 of models trained specifically on human faces (0.390 mAP@0.50). In contrast, human-face detectors showed dramatic performance degradation when applied to primate faces, achieving only 0.585 mAP on our PrimateFace test set compared to 0.775 mAP for PrimateFace-trained detectors (**Fig. 3c**). This disparity demonstrates that PrimateFace’s inherent morphological heterogeneity provides a richer training foundation that transfers effectively across species, while human-centric training captures only a narrow slice of possible facial variation, suggesting cross-primate variation may provide natural data augmentation that enhances generalization.

### PrimateFace enables seamless landmark conversion

Researchers working across primate facial analysis often encounter a fundamental interoperability challenge: existing datasets, pre-trained models, and analysis pipelines employ different landmark conventions (**Fig. 4a**), creating friction when integrating tools or comparing results across studies. A researcher might have DeepLabCut-MacaqueFace-55-landmarks annotations for their macaque data, but want to leverage human facial analysis tools expecting 68-keypoint inputs, or need to compare findings across datasets using incompatible landmark schemes. This fragmentation forces researchers to choose between specialized NHP tools and broader computer vision approaches^36^, limiting methodological flexibility.

To address this bottleneck, we developed a landmark conversion model – an attention-based multi-layer perceptron – that enables smooth conversion between the standard 68-keypoint configuration and our custom 48-keypoint scheme optimized for primate facial morphology (**Fig. 4b**, **Extended Data Fig. 6**). This conversion capability allows researchers to maintain their preferred analysis frameworks, while accessing the full ecosystem of facial analysis tools (e.g., see *Scientific Application 2* for an example of integration with InsightFace^8^), regardless of their original landmark format.

Our landmark conversion model achieved robust performance when converting from 68-keypoint to 48-keypoint configurations, with a mean per-joint position error (MPJPE) of 3.35 pixels on held-out 4K-resolution test data – equivalent to sub-pixel to 1.5-pixel error at standard 1080p resolution (**Fig. 4c**, **Extended Data Fig. 6**). Error distribution remained consistent across landmarks, ranging from 2.07 pixels for the nose bridge center (index 38) to 5.59 pixels for right brow outline landmarks (index 2). Qualitative assessment demonstrates faithful preservation of anatomical correspondence across diverse primate faces and poses (**Fig. 4d, Extended Data Fig. 6d**).

High landmark conversion accuracy enables practical workflow flexibility: researchers can now apply 68-keypoint human facial analysis models to NHP data, compare findings across datasets with different landmark schemes, and integrate specialized primate tools into broader computer vision pipelines without manual re-annotation. Moreover, researchers aren’t limited to working with the dense set of keypoints and can instead select keypoint subsets (e.g., perioral keypoints) with minimal fine-tuning in PrimateFace, enabling rapid yet robust analysis tailored to specific research questions. The lightweight architecture enables straightforward integration into existing workflows, facilitating cross-dataset comparisons that were previously incompatible. We provide workflows for fine-tuning, training from scratch, or training additional models such as graph neural networks, for landmark conversion.

### PrimateFace facilitates species-agnostic, accurate facial landmark estimation

Facial landmark localization enables detailed analysis of primate expressions, gaze patterns, and behavioral kinematics by providing consistent reference points that can be tracked over time in video sequences. When applied frame-by-frame to videos, these landmarks quantify dynamic facial movements – such as mouth opening during vocalizations, eye gaze direction, or brow movements during social interactions – providing objective kinematic measurements for behavioral analysis that previously relied on subjective manual assessment^6^. While existing NHP landmark models like DeepLabCut’s MacaqueFace represent pioneering efforts for individual species, no solution has achieved reliable landmark estimation spanning the full primate order. PrimateFace-trained models address this limitation, delivering robust 68-keypoint facial landmark estimation across all six primate superfamilies using top-down pose estimation architectures that build upon face detection outputs (**Fig. 5a,b**). We trained multiple state-of-the-art pose estimation models on the PrimateFace dataset using default hyperparameters, interoperable with platforms including MMPose, Ultralytics, DeepLabCut, and SLEAP for seamless integration (**Fig. 5d**).

#### PrimateFace landmarks outperform human and species-specific models across primate faces

To evaluate landmark estimation performance, we compared PrimateFace-trained models against human facial landmark models and existing primate-specific approaches. Our models employ a top-down pose estimation framework that first detects faces and then estimates landmarks within bounding boxes, enabling flexible integration with other frameworks (e.g., Ultralytics or mmdetection). For comparison, we evaluated human-face landmark models and DeepLabCut’s species-specific MacaqueFace model (**Fig. 5a**). While MacaqueFace uses a bottom-up approach that directly detects keypoints, we provided cropped face bounding boxes to optimize its performance, effectively creating similar input conditions. Across frameworks, top-down models trained on PrimateFace data perform comparably when using consistent training protocols (**Fig. 5d**).

Qualitative comparison of landmark estimation approaches reveals the advantages of cross-species training (**Fig. 5a**). PrimateFace models consistently localize facial landmarks across diverse primate genera, while human-face models show systematic failure modes – particularly misinterpreting mouth contours as facial boundaries in Cercopithecoidea, leading to cascading errors in nose and mouth landmark placement (see **Fig. 5a**, 2nd row, 5th column). Human models completely fail on morphologically distant species like Lemuroidea, as seen with Daubentonia. The DeepLabCut MacaqueFace-55 model occasionally performs well (see **Fig. 5a** third row, Lorisoidea and Cercopithecoidea columns) but demonstrates the limitations of species-specific training when applied beyond its target domain.

#### Facial landmark estimation maintains consistent performance across primate superfamilies

Quantitative evaluation across primate superfamilies revealed performance patterns reflecting both data availability and morphological constraints. Models exhibited lowest performance on Tarsioidea due to limited training data and their distinctive facial morphology – characterized by proportionally enormous eyes and compressed facial features that differ substantially from other primates. PrimateFace landmark estimators achieved optimal results on Hominoidea and Lemuroidea (**Fig. 5b**). Performance variation across genera (**Extended Data Fig. 2b**) reflects multiple factors: high-performing genera (Macaca, Pan) benefit from extensive datasets, both within and beyond the laboratory setting, and share more typical primate facial proportions, while challenging cases (Alouatta, the Howler Monkey) involve rapid facial movements during vocalizations that stress landmark tracking algorithms (see *Scientific Application 3*). This breakdown demonstrates the value of taxonomically-informed evaluation and highlights both data collection priorities and morphological considerations for future dataset expansions.

#### PrimateFace models maintain human facial landmark estimation performance

Cross-domain evaluation on COCO-WholeBody-Face demonstrated the generalization advantages of morphologically diverse training. PrimateFace-trained models achieved 0.061 normalized mean error (NME) on human landmark estimation – within 0.008 of human-face baselines (0.053 NME) (**Fig. 5c**). In contrast, human-face models showed substantial degradation on primate faces (0.122 vs. 0.029 NME on PrimateFace data, Δ 0.093). This reinforces that cross-species training provides feature representations that transfer effectively to humans, while human-centric training fails to capture broader facial variation.

This holds implications for comparative primatology and translational research. Researchers can now apply consistent landmark schemes across species for evolutionary studies, employ unified analysis pipelines for mixed human-NHP datasets, and leverage the growing ecosystem of human facial analysis tools for non-human primate behavioral research regardless of existing computational infrastructure.

### PrimateFace accelerates and enables diverse research applications

The PrimateFace dataset and models provide a foundation for diverse scientific investigations across computational primatology. To demonstrate their utility, we present five applications spanning face presence logging, individual recognition, vocal-motor coupling analysis, social attention dynamics, and unsupervised behavior discovery, each accompanied by documented workflows and code notebooks for immediate replication. These examples showcase how PrimateFace enables rapid development of sophisticated analytical pipelines that address distinct challenges in understanding primate behavior, while also demonstrating cross-species generalization capabilities that extend to human applications.

#### Scientific Application 1: Automated Time-Stamping Pipeline

Manually logging when individuals are present in video recordings is a fundamental but exceptionally time-consuming task in observational research. PrimateFace cross-species face detectors provide the foundation for automating this critical step. We demonstrate an automated time-stamping pipeline that leverages a PrimateFace face detector to process video footage frame-by-frame (**Fig. 6a**). This effectively compresses days or weeks of raw video into concise, interpretable visualizations that summarize individual visibility over time. The pipeline exports to Behavioral Observation Research Interactive Software (BORIS) format, enabling researchers to load footage with time-stamped ‘Face Present’ event states into this widely-used ethological annotation platform. Presence–absence rasters constitute the detection histories used in occupancy analysis – an approach first developed for camera-trap surveys to infer site use and animal movement^37^. This automated visibility timeline significantly enhances the efficiency of longitudinal monitoring, enabling researchers to quickly identify patterns of individual presence, co-occurrence with conspecifics, solitary periods, and interactions with the environment.

**Figure 6.**
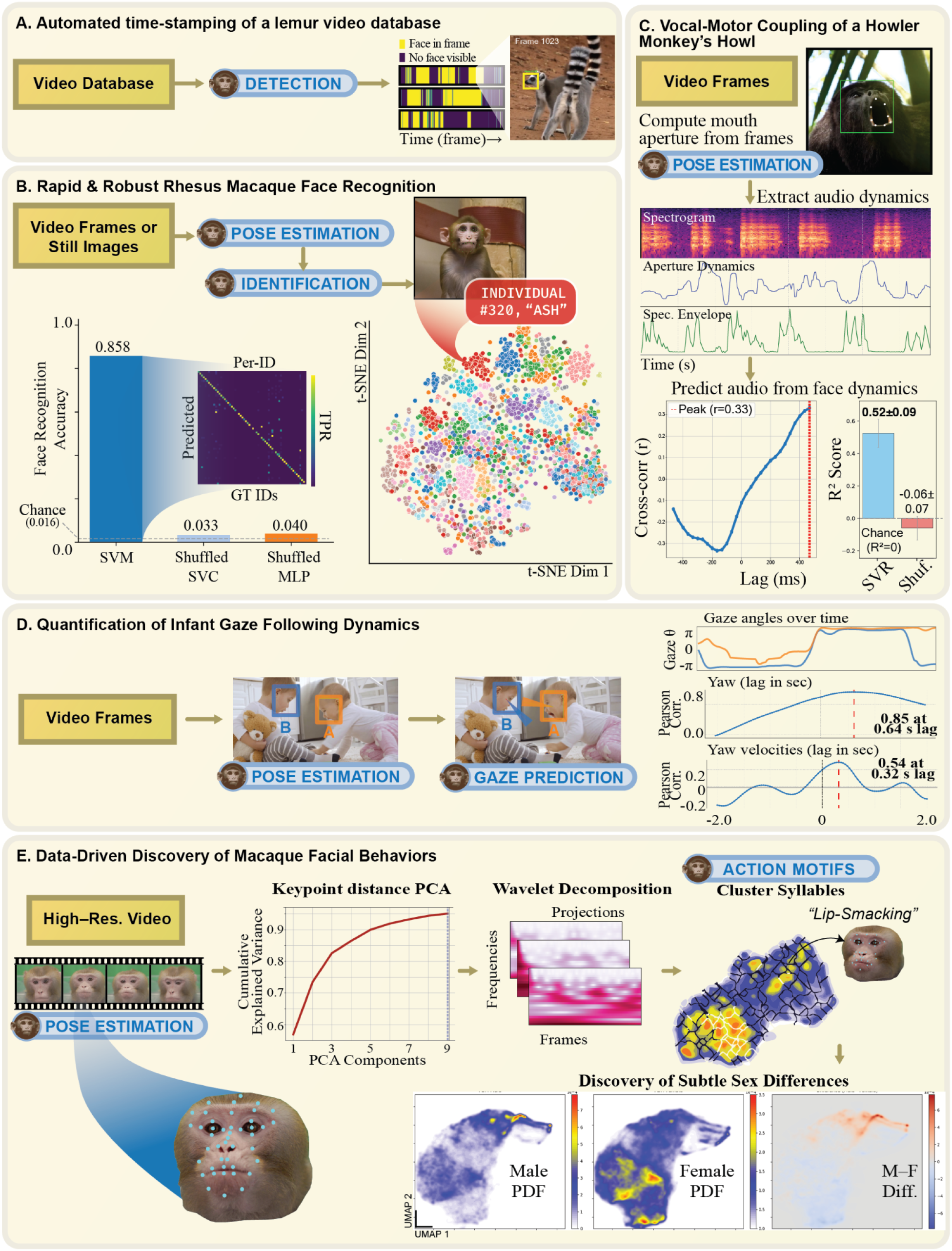
PrimateFace accelerates and enables diverse research applications. **A.** Automated time-stamping of lemur faces in video yields presence-absence raster plots, compressing days of footage into minutes of review. **B.** PrimateFace enables researchers to train personalized face recognition classifiers in under an hour, achieving 86% accuracy across 62 individuals. Inset: confusion matrix comparing predicted versus ground-truth (GT) identities. Right: Unsupervised clustering of face ID embeddings reveals discrete individual-specific clusters. **C.** Extraction of facial kinematics and audio during howler monkey vocalizations reveals cross-modal coupling through time-aligned acoustic (spectrogram) and kinematic (mouth aperture) analysis. **D.** Gaze-following dynamics among human infants estimated from head pose, showing correlated gaze directions and rotational velocities. **E.** High-resolution macaque facial behavior analysis via pose unsupervised action discovery reveals subtle yet discrete facial movement (e.g., left-turning or more stereotyped movements like lip-smacking) with fine-grained male-female differences captured through density analysis. PrimateFace transforms raw pixels into interpretable, high-resolution behavioral events. **Acronyms:** SVM/C, support vector machine/classifier; MLP, multi-layer perception; Dim, dimension; Spec., spectrogram; SVR, support vector regression; Res., resolution; PCA, principal component analysis; PDF, probability density function.

#### Scientific Application 2: Rapid Face Recognition of 62 Individual Monkeys

While researchers have previously developed automated individual recognition systems for specific species and datasets, these efforts require considerable technical expertise, species-specific data, and extensive feature engineering. Previously, researchers had to hand-crop every image (or by first training a customized face detection and alignment model) before identification could even start. PrimateFace automates these labor-intensive front-end steps in a single forward pass, freeing effort for experimental design rather than data wrangling (**Fig. 6b**).

To demonstrate this capability, we applied PrimateFace’s off-the-shelf face detection and 68-keypoint facial landmark estimation models to a public dataset of rhesus macaque faces^38^, comprising 62 individuals after filtering for sufficient samples. We leveraged InsightFace^8^, a popular human face analysis library implementing the ArcFace recognition model, to generate 512-dimensional embeddings from aligned primate faces.

The streamlined pipeline involved: 1) detecting faces with PrimateFace, 2) extracting facial landmarks with PrimateFace landmark estimator, 3) aligning faces using the 5-landmark convention (see **Fig. 4a** far left), 4) generating ArcFace embeddings from the aligned faces, and 5) training a support vector machine (SVM) classifier on these embeddings. This entire process, from raw images to a trained recognition model, can be executed in an hour.

The resulting classifier achieved robust top-1 identification accuracy of 0.858 across the 62 individuals – significantly above chance levels (1/62 ≈ 0.016) and shuffled-label controls (**Fig. 6b**). The confusion matrix reveals strong recognition performance, while t-SNE visualization of ArcFace embeddings show monkey identity-based clustering (**Fig. 6b right**). This demonstrates how PrimateFace eliminates barriers to individual recognition, enabling researchers to rapidly create accurate ID systems without specialized development.

#### Scientific Application 3: Fine-Grained Vocal-Motor Coupling Analysis

Understanding the coordination between facial movements and vocal production is crucial for deciphering the mechanisms and communicative intent of vocal signals across species^39,40^, yet precisely tracking mouth dynamics during active vocalizations has remained exceptionally challenging. Open mouth configurations – critical during calls, howls, and other vocalizations – are notoriously difficult to landmark accurately due to rapidly changing apertures, occlusions, and species-specific oral morphologies.

PrimateFace overcomes this longstanding limitation by providing robust landmark estimation even during dynamic vocalizations. To demonstrate this capability, we applied PrimateFace detection and landmark models to videos of howling howler monkeys. We first fine-tuned the landmark estimation model on howler-monkey-specific data for enhanced mouth tracking performance, and then extracted precise mouth aperture dynamics throughout the vocalization sequence (**Fig. 6c**). Cross-correlation analysis revealed modest coupling (r = 0.33) between mouth aperture and vocal envelope. More importantly, a support vector regression (SVR) model successfully predicted acoustic dynamics from facial kinematics, significantly outperforming shuffle controls (**Fig. 6c**).

This workflow opens new possibilities for addressing questions that were difficult to tackle systematically. By automating vocal-motor analysis, PrimateFace makes it practical to examine vocal repertoires across species, investigate individual differences in coordination patterns, and compare vocal mechanisms across the primate phylogeny at scales that can inform our understanding of communication system evolution and neural control mechanisms.

#### Scientific Application 4: Quantifying Gaze Following in Human Infants

The ability to follow another’s gaze is a cornerstone of social cognition, crucial for joint attention and understanding communicative intent^41^. However, manually coding gaze dynamics and behavioral synchrony in naturalistic interactions – whether studying human infants or non-human primates – is extremely labor-intensive and subject to observer bias, severely limiting the scale of developmental studies possible. Automated analysis requires precise face detection combined with gaze estimation models – specialized computer vision systems that predict where someone is looking (i.e., the angular direction) based on eye and head position.

PrimateFace’s generalizable cross-species face detection provides the essential foundation for integrating with such specialized models. We combined PrimateFace detection with a state-of-the-art gaze estimation model known as GazeLLE^42^ on Adobe Stock footage of two interacting human infants (**Fig. 6d**). PrimateFace first localized each face and maintained relative identities with temporal tracking, then GazeLLE estimated gaze direction for each infant over time. Cross-correlation analysis across lags from −2 to +2 seconds revealed behavioral synchrony: yaw angles exhibited Pearson correlation of 0.85 at 0.64 s lag, while yaw velocities showed correlation of 0.54 at 0.32 s lag – critical markers of social engagement that have been difficult to quantify objectively at scale.

By combining PrimateFace with specialized visual models, researchers can objectively measure behavioral synchrony across larger sample sizes and longer time periods than manual coding allows. This enables systematic investigation of attention coordination emergence, caregiver-infant synchrony patterns, and individual differences in social attention coupling.

#### Scientific Application 5: Data-Driven Discovery of Subtle Facial Movements

Primate facial communication involves a rich repertoire of dynamic movements, yet traditional ethological approaches rely on a priori manually defined behavioral categories that may miss subtle patterns or introduce observer bias. The challenge of systematically discovering the full behavioral repertoire – including rare, transitional, or species-specific movements – has remained a fundamental bottleneck in understanding primate communication.

Data-driven identification of ‘behavioral syllables’ – brief, reusable movement patterns automatically extracted from continuous video – has emerged as a powerful framework in behavioral science, enabling objective cross-animal comparisons by decomposing complex behavior into discrete, interpretable units that reveal hidden structure in natural behavior (e.g., MotionMapper^43^ or Keypoint-MoSeq^44^).

PrimateFace’s precise facial landmark tracking enables applying this paradigm to primate facial communication. We recorded facial interactions between pairs of chaired macaque monkeys across experimental sessions using high-frame-rate (120 FPS) and high-resolution (4K) cameras over 15-minute sessions. We utilized PrimateFace’s 48-keypoint estimation model to extract detailed facial kinematics through a comprehensive pipeline: video sequences underwent face detection and pose estimation, followed by computation of inter-landmark distances and velocities. Principal component analysis retained the top 9 components capturing the majority of facial movement variance, which were then processed through a modified MotionMapper pipeline with wavelet decomposition, UMAP embedding, and watershed clustering to identify over 80 recurrent movement patterns (**Fig. 6e**). Expert validation confirmed that discovered patterns corresponded to both stereotyped facial movements (lip-smacking) and subtle behaviors (head turns, eye-blinking). Notably, this data-driven approach revealed sex-specific differences in facial communication repertoires – subtle patterns invisible to human experimenters during data collection. This workflow enables objective discovery of species-specific behavioral vocabularies and direct linking of facial movements to neural activity recordings^31^.

## Discussion

PrimateFace represents a significant advancement from manual coding to high-throughput quantitative analysis of primate facial behavior. We provide, to our knowledge, the first publicly available dataset of this scale and taxonomic diversity for primates, with performant models tailored for cross-species analysis. Our standardized dataset and robust models empower researchers to analyze previously intractable volumes of video data with enhanced speed and precision. The cross-species training approach delivers models that generalize effectively across the primate order while maintaining competitive performance on human face benchmarks, demonstrating the advantages of morphologically diverse training data. PrimateFace offers transformative potential for advancing primate research across social behavior, cognition, ecology, welfare, and evolution.

The five scientific applications showcase PrimateFace’s versatility: automated visibility time-stamping compresses days of footage into rapid review; individual recognition systems can be trained in under an hour; vocal-motor coupling analysis reveals cross-modal coordination during vocalizations; gaze dynamics quantify social attention patterns; and unsupervised behavioral discovery identifies both stereotyped and subtle facial movements, including sex-specific differences invisible to human observers. Beyond social interaction research^45^, PrimateFace enables analysis of chewing dynamics^46^, pain assessment^32^, and other non-social behaviors. The benefits of studying movement across species extend beyond primatology, fostering novel applications in neuroscience^47,48^, medicine^49^, and machine learning^50^.

Despite its breadth, PrimateFace does not exhaustively cover all primate species, and rarer genera remain underrepresented. Our models can encounter difficulties with extreme occlusions, lighting, or unusual viewpoints, particularly for less represented species. Current annotations focus on face bounding boxes and landmarks rather than discrete facial action units (AUs) or 3D structure. Automated cross-species AU detection remains challenging due to species-specific musculature, fur patterns, and limited annotated training data^5,51^. Future directions include strategic dataset expansion for underrepresented taxa and behavior, incorporating facial action units, and integrating novel deep learning models.

PrimateFace positions the field to drive progress across multiple domains. Within computational primatology, PrimateFace enables training foundation models for diverse visual tasks from expression analysis to behavioral prediction. For computer vision, PrimateFace provides challenging benchmarks for few-shot learning and cross-species domain adaptation. Broader implications include enhanced primate welfare monitoring, improved conservation efforts through individual tracking, and advances in comparative psychology and neuroscience. By integrating deep learning with comprehensive, open datasets, PrimateFace demonstrates the transformative potential of machine learning applied to complex biological questions. We openly release the PrimateFace dataset, models, and software to the community, inviting contributions to continually expand this resource and advance primate research.

## Methods

### Data Sources and Permissions

The PrimateFace dataset contains over 260,000 primate images spanning humans and non-human primates. Approximately 200,000 images were aggregated and re-annotated from publicly available scientific datasets and repositories (**Extended Data Table 1**), utilized in accordance with their respective terms of use and licensing. An additional ∼60,000 images were obtained through targeted web searches of publicly accessible primate imagery. Web collection excluded copyrighted material not explicitly licensed for reuse.

### Newly Collected Non-Human Primate Data

The novel collection of 60,000 non-human primate images, designed to enhance taxonomic balance within PrimateFace, was sourced from existing internal laboratory archives (where ethical approvals for original data collection were already in place), from public domain sources, and from collaborator contributions (for which they held relevant IACUC approvals). Specifically, the high-resolution video data of rhesus macaques (*Macaca mulatta*) used for the behavioral syllable discovery demonstration in **Fig. 6e** were collected at the University of Pennsylvania. These procedures were approved by the Institutional Animal Care and Use Committee (IACUC) of the University of Pennsylvania and were performed in accordance with all relevant institutional and national guidelines and regulations for the ethical treatment of animals.

### Use of Human Imagery for Model Benchmarking and Demonstrations

While the openly released PrimateFace dataset focuses on non-human primates, human face data was utilized in specific contexts to demonstrate cross-species model generalization. Our models were benchmarked against standard human face datasets (WIDERFace^9^, COCO-WholeBody-Face^10^) – established computer vision benchmarks sourced from public repositories where ethical considerations and consent were managed by the original data creators (see **Extended Data Table 1**). Additionally, the gaze-following demonstration utilized video of human infants obtained from Adobe stock footage, with no direct human subject involvement. No human images are included in the distributed PrimateFace dataset.

### Data Release

The PrimateFace dataset comprises re-annotated images from existing publicly available research datasets and public domain sources (see **Extended Data Table 1**). The dataset is intended to advance understanding of primate behavior and support non-invasive research methodologies that reduce the need for additional animal studies.

### PrimateFace Dataset Construction

All images underwent re-annotation to conform to PrimateFace’s standardized labeling protocols. Additional images were acquired through systematic web searches targeting publicly accessible content from academic institutions, research databases, and zoological society websites. To address taxonomic underrepresentation across the primate order, approximately 60,000 additional images were curated to achieve balanced representation across 60+ primate genera. This curation process set targets of at least 700 images per genus, with particular emphasis on prosimian and New World monkey taxa that are typically underrepresented in computer vision datasets. Image selection prioritized clear facial views, diverse individuals, and varied environmental contexts to maximize training data utility. The resulting PrimateFace dataset encompasses 260,000+ images representing 60+ primate genera. The collection captures diverse real-world conditions including natural habitats, zoological settings, and research contexts from previously published studies, with varied lighting, age ranges from infants to adults, multiple expressions, and social interactions (**Fig. 2, Suppl Fig. 1**). This comprehensive scope provides a robust foundation for cross-species facial analysis.

### Data-Annotation Pipeline

#### DINOv2-Guided Core Set Selection for Initial Annotation

To bootstrap annotation with maximum visual diversity, we developed a data-driven selection pipeline for choosing 10,000 initial training images from over 200,000 ‘starting’ images (**Extended Data Table 1**, **Extended Data Fig 3b**). We computed 768-dimensional DINOv2 ViT-Base embeddings, then employed hybrid clustering and Farthest Point Sampling to maximize both broad manifold coverage and local diversity within clusters.

#### Sampling Efficiency Benchmark for Detector Training

We systematically compared DINOv2-guided sampling against random selection for face detector training across annotation budgets (500-8000 images). Training subsets were stratified using PCA and k-means clustering in embedding space, with identical architectures evaluated on held-out test data (**Fig. 3d**).

#### Embedding-space visualization

To qualitatively assess the learned DINOv2 representations, we created two complementary visualizations. We projected the 768-dimensional CLS embeddings to two dimensions using UMAP and colored points by 100-way k-means clustering (**Extended Data Fig. 2b**). We also examined model attention patterns by processing exemplar images through the ViT-Base network, applying PCA to patch tokens to create pseudo-RGB mosaics and reshaping CLS-token attention scores from all 12 transformer layers onto the spatial patch grid (**Extended Data Fig. 4**). These visualizations reveal clustering patterns in feature space and which image regions the model prioritizes during feature extraction.

#### Face Bounding Box Annotation

All primate faces in the PrimateFace dataset required standardized bounding box annotation across diverse taxa. For images with existing sparse keypoint annotations, initial face bounding boxes were computationally derived to encompass these features. For remaining images, including the newly collected set, bounding boxes were generated through our semi-automated pipeline. Our annotation guidelines defined face regions as requiring at least two of three key features to be clearly visible: eyes, nose, and mouth, with boxes drawn to tightly enclose the facial region while excluding ears. This standardized approach maintains annotation reliability across species with varying ear morphology and visibility.

#### Facial Landmark Annotation Schemas

PrimateFace provides annotations for two distinct facial landmark configurations, selected to maximize utility for both primate-specific research and computer vision applications. Facial landmark schemes range from sparse 5-keypoint configurations for basic face alignment, to the 55-keypoint DeepLabCut MacaqueFace model for macaque analysis, to the standard 68-keypoint schema for human face analysis (**Fig. 4a**).

The first PrimateFace schema is a 48-keypoint configuration developed for cross-species analysis in non-human primates. This schema densely maps key facial features including eyes, nose, mouth, and jawline. The design ensures anatomical relevance and consistent identifiability across diverse NHP morphologies. The second is the standard 68-keypoint schema, including detailed outlines of eyebrows, eyes, nose, mouth, and jaw. This configuration facilitates interoperability with existing human-centric models and datasets, enabling comparative studies and leveraging existing human facial analysis tools.

#### Iterative Pseudo-Labeling and GUI-Assisted Correction

To manage large-scale annotation efficiently, we implemented an iterative, semi-automated workflow (**Fig. 1**). Following manual annotation of the DINOv2-selected core set, initial models for face detection and landmark estimation were trained to generate pseudo-labels on new image batches. These machine-generated annotations were reviewed and corrected by human annotators using a custom-developed graphical user interface (**Extended Data Fig. 3c**).

The GUI presents proposed bounding boxes and landmark configurations overlaid on images, enabling annotators to rapidly accept, reject, or adjust annotations. Completed annotations were saved in COCO-JSON format. This iterative cycle of training models, generating pseudo-labels, and human correction continued until adequate performance across primate genera.

#### Quality Control

Rigorous measures ensured annotation accuracy and consistency throughout the process. All annotations were performed or reviewed by at least two expert reviewers. A subset employed double-pass annotation with discrepancies resolved through senior review.

Inter-annotator agreement was periodically assessed using Intersection over Union (IoU) for bounding boxes, with annotations below 0.85 IoU triggering consensus review. Facial landmark accuracy was evaluated using Normalized Mean Error (NME) between annotators. Regular calibration sessions maintained high inter-rater reliability.

#### Semi-automated genus classification

To enable taxonomic balancing across 60+ primate genera, we developed a semi-automated classification pipeline. Face crops from PrimateFace detection models were submitted to Gemini 2.5 Pro API for genus classification. VLM predictions were integrated into our annotation GUI (**Extended Data Fig. 3c-d**), enabling annotators to rapidly accept correct classifications or provide corrections. This prioritized collection from underrepresented taxa to achieve target allocations of ∼500-600 images per genus, ensuring balanced representation across the primate order (**Extended Data Fig. 1**). A subset underwent double-pass validation by taxonomic experts. This pipeline accelerated curation while preserving essential taxonomic expertise.

##### PrimateFace Model Development

###### Overview of Model Training Approach

We developed deep learning models for automated facial analysis across primate species, adapting state-of-the-art architectures for primate morphological diversity. Multiple frameworks were used to maximize research accessibility: MMDetection and Ultralytics for detection, MMPose, DeepLabCut, and SLEAP for landmark estimation. All models used *default training parameters* for reproducibility, with training-validation-test splits stratified by genera to assess cross-species generalization.

###### Face Detection Models

We fine-tuned several face detection models, including Cascade R-CNN and RetinaNet architectures using MMDetection, initialized with COCO pretrained weights. YOLOv8 models were trained using Ultralytics for real-time applications. Standard inference protocols used confidence thresholding (0.3-0.5) and Non-Maximum Suppression.

###### Facial Landmark Estimation Models

We trained several facial landmark estimation models. HRNet-W32 demonstrated optimal performance for both 68-keypoint and 48-keypoint configurations using MMPose, initialized with COCO-WholeBody pretrained weights. Additional models were trained across Ultralytics, DeepLabCut, and SLEAP frameworks and were evaluated using Normalized Mean Error (NME) metrics (**Fig. 5d**).

###### Landmark Configuration Conversion Model

To enable seamless conversion between 68-keypoint and 48-keypoint landmark configurations, we developed an attention-enhanced multi-layer perceptron for coordinate regression. The architecture uses self-attention to capture global keypoint relationships before regressing target coordinates (**Extended Data Fig. 6**).

Ground-truth data consisted of 68-keypoint annotations following standard conventions, with 48-keypoint targets derived using the pretrained DeepLabCut MacaqueFace-55 model. To ensure representative sampling across poses and identities, we applied k-means clustering to PCA-reduced keypoint coordinates before stratifying into training/validation/test sets (70/15/15). Separate bidirectional conversion models were trained using AdamW optimization, minimizing mean squared error between predicted and target coordinates. Performance was evaluated using Mean Per-Joint Position Error (MPJPE).

###### Qualitative Comparisons

We compared our PrimateFace-trained Cascade R-CNN model against an off-the-shelf human face detection model and GroundingDINO’s zero-shot detection capabilities. The first two models were implemented via MMDetection and confidence-thresholded at 0.75, while GroundingDINO was accessed through the Roboflow library, prompted with “face,” and thresholded at 0.5 due to its open-vocabulary nature requiring lower confidence thresholds. Bounding box predictions are visualized in yellow (**Fig. 3a**).

We compared facial landmark estimation HRNetV2-W18 models trained on PrimateFace data (top row) versus COCO-WholeBody-Face (middle row) (**Fig. 5a**). The COCO-WholeBody-Face model used pretrained MMPose checkpoints without retraining. We additionally evaluated the DeepLabCut MacaqueFace model zoo checkpoint for comparison, which uses a distinct 55-keypoint configuration. All landmark models were confidence-thresholded at 0.3.

### Evaluation Metrics

Face detection model performance was evaluated using mean Average Precision (mAP) following standard object detection benchmarks. We primarily report mAP@0.5, which calculates Average Precision at an Intersection over Union (IoU) threshold of 0.5. For the DINOv2 sampling efficiency comparison (**Fig. 3d**) and genus-level performance analysis (**Extended Data Fig. 2a**), we used mAP@0.50:0.95, averaging AP across IoU thresholds from 0.5 to 0.95 with 0.05 step increments. Average Precision is calculated as the area under the precision-recall curve, where precision is the ratio of true positives to total detections (TP/(TP+FP)) and recall is the ratio of true positives to total ground truth instances (TP/(TP+FN)). A detection was considered a true positive if its IoU with a ground truth bounding box exceeded the threshold and the ground truth had not been previously matched.

For facial-landmark models we report the Normalized Mean Error (NME). For an image with *L* landmarks, NME is the average Euclidean distance between each predicted landmark (**x**_pred,*i*_, **y***_pred,i_*) and its ground-truth counterpart (**x**_gt,*i*_, **y***_gt,i_*), normalized by the inter-ocular distance *D* (the distance between the outer eye corners):

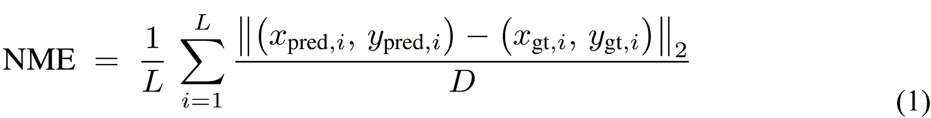

The dataset-level NME is the mean across all test images. Lower NME values indicate better landmark localization accuracy.

### Benchmarking Details

#### PrimateFace Dataset Splits and Evaluation Protocol

For model training and evaluation, PrimateFace data was partitioned into training (17,968 images; 17,397 bboxes; 10,827 images with keypoints), validation (996 images; 1,362 bboxes; 741 images with keypoints), and test (996 images; 1,311 bboxes; 743 images with keypoints) sets using stratification by primate genera. This ensures robust generalization assessment to potentially unseen or underrepresented genera during training. All reported PrimateFace performance metrics were computed on the held-out test set.

#### Human Dataset Evaluation Protocols

To assess cross-species generalization, we conducted bidirectional evaluation between PrimateFace and standard human face benchmarks following MMDetection (mmdetection/[WIDERFace](https://github.com/open-mmlab/mmdetection/tree/cfd5d3a985b0249de009b67d04f37263e11cdf3d/configs/wider_face)) and MMPose (mmpose/[COCO-WholeBody-Face](https://github.com/open-mmlab/mmpose/blob/main/docs/en/dataset_zoo/2d_face_keypoint.md)) documentation protocols.

For face detection, we used the WIDERFace dataset (12,880 training images; 3,226 validation images) and trained RetinaNet models on both PrimateFace and WIDERFace training sets. Cross-domain evaluation assessed PrimateFace-trained models on WIDERFace validation data and PrimateFace test data.

For facial landmark estimation, we used the COCO-WholeBody-Face dataset (118,582 training images; 5,000 validation images) and trained HRNet-W18-Dark models on both PrimateFace and COCO-WholeBody training sets. Cross-domain evaluation followed standard COCO keypoint evaluation protocols, with NME normalized by inter-ocular distance as defined for human faces.

Note that both WIDERFace and COCO-WholeBody-Face provide validation sets rather than test sets for public evaluation, which we used for all reported human benchmark results.

### Application Demonstration Setups

#### Automated Time-Stamping Pipeline

Temporal analysis of individual visibility is fundamental for behavioral studies requiring precise event timing. Using lemur footage from YouTube Research API (not included in PrimateFace distribution), video sequences were processed frame-by-frame using PrimateFace-trained Cascade R-CNN face detection. Face detections were confidence-thresholded and filtered using Non-Maximum Suppression, with temporal presence data generating raster plots visualizing individual visibility over time (**Fig. 6a**). This demonstrates automated behavioral time-stamping capabilities across diverse primate species and video conditions.

#### Macaque Face Recognition

Rapid individual identification is critical for longitudinal behavioral studies and population monitoring. Using the Witham & Bethell (2019) rhesus macaque dataset (62 individuals with ≥3 samples; not included in PrimateFace), we implemented a five-step pipeline: face detection, 68-point landmark extraction, face alignment to 112×112 pixels, ArcFace feature embedding extraction (512-dimensional), and classifier training with 70/30 train/test splits stratified by identity. Support Vector Machine and Multi-Layer Perceptron classifiers achieved high accuracy with optimized hyperparameters, evaluated using confusion matrices and t-SNE visualization (**Fig. 6b**). This demonstrates how PrimateFace enables rapid development of species-specific recognition systems.

#### Howler Monkey Vocal-Motor Coupling Analysis

Understanding the coordination between facial movements and vocalizations provides insights into primate communication mechanisms and evolutionary origins of speech. Using howler monkey footage from YouTube Research API (not included in PrimateFace distribution), we analyzed temporal relationships between mouth kinematics and acoustic output (**Fig. 6c**).

PrimateFace models detected faces and estimated 68 facial landmarks per frame, with landmarks aligned to standard coordinate space to minimize head motion artifacts. Frame-by-frame mouth aperture was calculated as the average Euclidean distance between three pairs of corresponding inner-lip landmarks, with the time-series undergoing median filtering, detrending, and normalization. Audio tracks were processed into log-magnitude spectrograms with temporal envelopes derived from maximum magnitude across frequency bins. Cross-correlation analysis assessed temporal relationships over ±500 ms lag ranges, while Support Vector Regression models predicted audio envelope dynamics from facial kinematic features. This demonstrates PrimateFace’s utility for quantifying vocal-motor coordination, enabling automated analysis of communication behaviors across primate species without manual landmark annotation.

#### Infant Gaze-Following

Joint attention and gaze coordination are fundamental markers of early social cognition development, requiring precise temporal analysis of multi-individual interactions. Using stock footage of interacting human infants from Adobe Stock (used for training, not distribution), we demonstrated automated quantification of gaze dynamics and behavioral synchrony (**Fig. 6d**).

PrimateFace-trained Cascade R-CNN detected faces per frame with IoU-based tracking maintaining consistent identities. Gaze direction and confidence were estimated using GazeLLE models, with continuous gaze trajectories smoothed using low-pass Butterworth filtering. Yaw velocity was computed as frame-to-frame derivatives of smoothed angles, with both measures z-score normalized for cross-individual comparison. Behavioral synchrony was quantified through cross-correlation analysis of yaw angle and velocity time series over ±2 second lag ranges, extracting peak correlation coefficients and temporal lags to characterize gaze coordination dynamics.

This demonstrates PrimateFace’s utility for objective quantification of joint attention patterns and social synchrony, enabling automated analysis of developmental and social behaviors that traditionally require labor-intensive manual coding.

#### Behavioral syllable discovery

Automated identification of discrete behavioral units enables objective quantification of communication repertoires and social dynamics without subjective manual coding. Using novel high-resolution (4K, 120 fps) recordings of dyadic facial interactions between rhesus macaques from Platt Labs at UPenn (used for training, not distribution), we demonstrated unsupervised behavioral syllable discovery (**Fig. 6e**).

PrimateFace models detected faces and extracted 48-keypoint landmark trajectories, with time-series computed from all pairwise Euclidean distances between keypoints for each subject. Principal Component Analysis applied to concatenated distance matrices obtained lower-dimensional facial kinematic representations, followed by continuous wavelet transform to capture time-frequency movement characteristics.

Following MotionMapper framework principles^43^, these features generated a reference behavioral embedding space, with UMAP dimensionality reduction and watershed segmentation identifying discrete regions corresponding to putative behavioral syllables. Probability density functions of syllable usage were computed separately for male and female macaques, with difference maps identifying sex-specific patterns in facial behavioral expression.

This demonstrates PrimateFace’s capability for unsupervised discovery of discrete communicative behaviors, enabling automated analysis of social repertoires and individual differences that would otherwise require extensive manual annotation by trained observers.

## Code and data availability

PrimateFace is freely available as open-source software, with documentation, guides, and notebook tutorials at https://github.com/KordingLab/PrimateFace.

## Acknowledgements

We would like to thank the labs of Michael L. Platt and Konrad P. Kording for their valuable advice and support. We also acknowledge the support of our funding agencies and institutions.

## Extended Data

**Extended Data Table 1.** PrimateFace Dataset Sources. A comprehensive curation of papers, datasets, and models at the intersection of machine learning and primatology can be found at https://github.com/KordingLab/awesome-computational-primatology. Any images extracted from open-source datasets were pulled from the ‘train’ split. HF indicates a dataset obtained from the Hugging Face ecosystem.

| Data Source | Images Extracted | Dominant Setting | Open-Source Face Model? |
| --- | --- | --- | --- |
| Animal Kingdom<sup>52</sup> | 981 | Wild | Body pose yes; Face no |
| AnimalFaceData<sup>26</sup> | 985 | Wild | No |
| AP-10k<sup>53</sup> | 1,074 | Wild | Body pose yes; Face no |
| APT-36k<sup>54</sup> | 6,960 | Wild | Body pose yes; Face no |
| Bristol2020<sup>55</sup> | 5,409 | Zoo | No |
| Web | 40,000 | Mix | No |
| Cayo Santiago Macaques | 42,574 | Wild | No |
| FairFace<sup>56</sup> | 4,971 | Mix | Human Face |
| Google OpenImages v7<sup>57</sup> | 2,022 | Mix | Body pose yes; Face no |
| HF ‘animal-dataset 1’ | 180 | Wild | Body pose yes; Face no |
| HF ‘Imagenet-monkey’ | 10,399 | Wild | No |
| MacaquePose<sup>13</sup> | 13,083 | Wild | Body pose yes; Face no |
| Adobe Stock | 500 | Stock | No |
| OpenMonkeyChallenge<sup>58</sup> | 80,000 | Wild | Body pose yes; Face no |
| OpenApePose<sup>59</sup> | 45,000 | Wild | Body pose yes; Face no |
| OpenMonkeyStudio<sup>12</sup> | 2,000 | Institution | Body pose yes; Face no |
| Philly Zoo (Personal Images) | 100 | Zoo | No |
| Platt Labs | 1,500 | Institution | No |
| Visiome<sup>60</sup> | 200 | Mix | No |
| WIDERFace<sup>9</sup> | 5,000 | Mix | Human Face (mmdetection) |
| COCO WholeBody-Face<sup>10</sup> | 5,000 | Mix | Human Face (mmpose) |
| Witham & Bethell 2019<sup>38</sup> | 3,100 | Institution | Yes (DeepLabCut) |
| **PrimateFace** | 271,038 |  | Yes (mmdet/pose, Ultralytics, DLC, SLEAP) |

**Extended Data Table 2.**
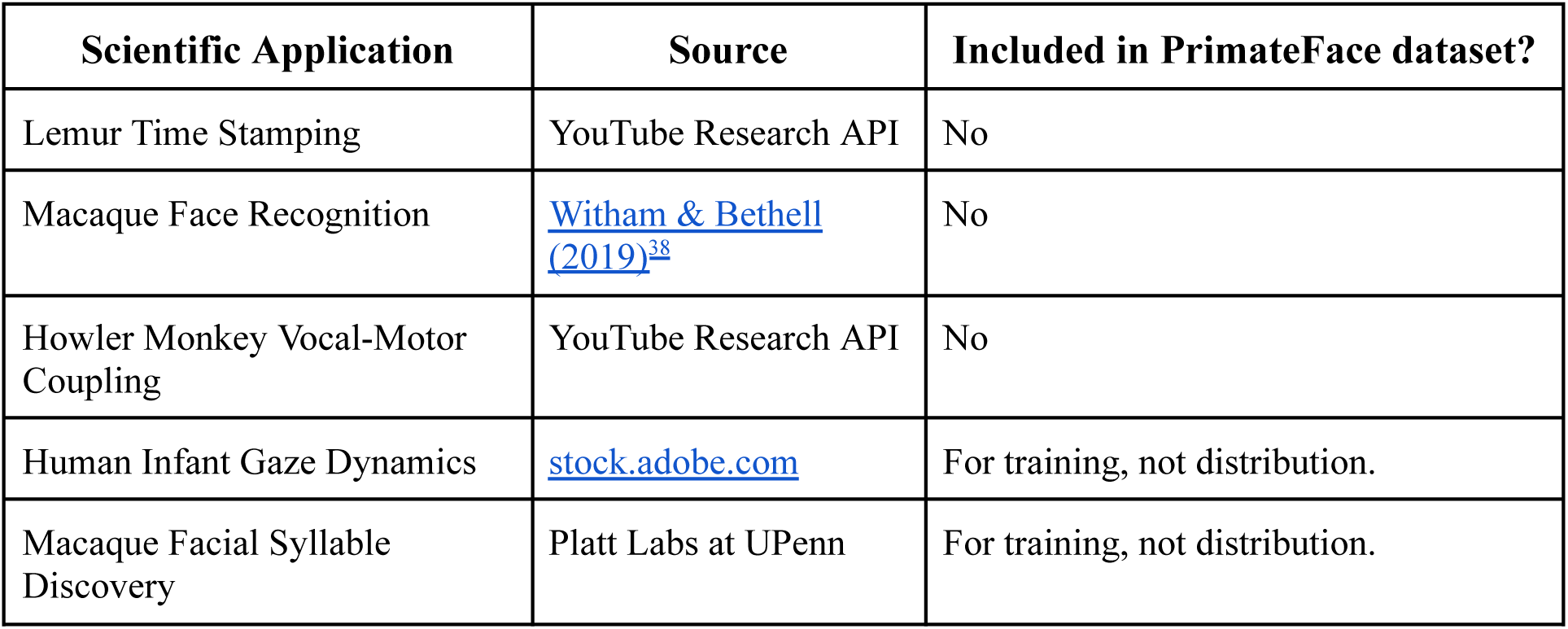
Data sources for scientific application demonstrations. Summary of datasets used to demonstrate PrimateFace’s utility across diverse research applications. Applications showcase the versatility of PrimateFace models across species (lemurs, macaques, howler monkeys), contexts (wild behavior, laboratory studies, developmental research), and analytical approaches (individual recognition, temporal analysis, unsupervised discovery). Data sources vary in availability and licensing: YouTube Research API data and published datasets remain separate from PrimateFace for licensing reasons, while stock photography and institutional data were used for model training but are not redistributed due to commercial or institutional restrictions.

**Extended Data Figure 1.**
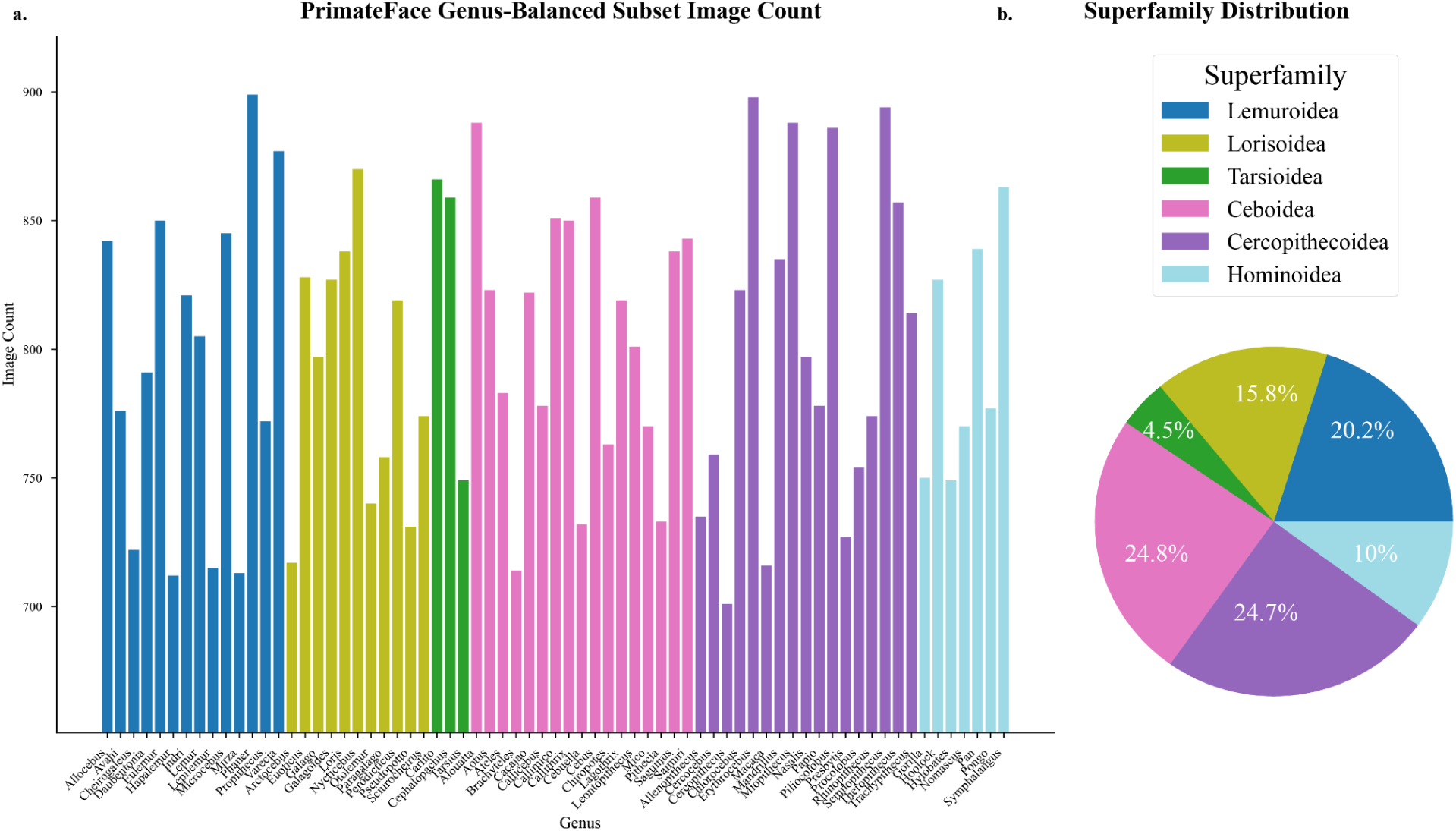
Taxonomic composition of PrimateFace’s balanced 60,000-image subset (v1). A. Bar plot showing image and annotation counts by genus within the strategically curated subset designed to enhance taxonomic balance across A. the primate superfamilies. This balanced composition ensures robust representation of morphologically diverse genera, including previously underrepresented groups such as tarsiers and prosimians, while maintaining sufficient sample sizes for effective model training across all six primate superfamilies. Genus-level balancing addresses the natural skew in available primate imagery toward more commonly photographed species, enabling development of truly cross-species facial analysis models. Note that this distribution reflects the initial v1 release and will evolve with future dataset expansions to incorporate community contributions and additional taxa.

**Extended Data Figure 2.**
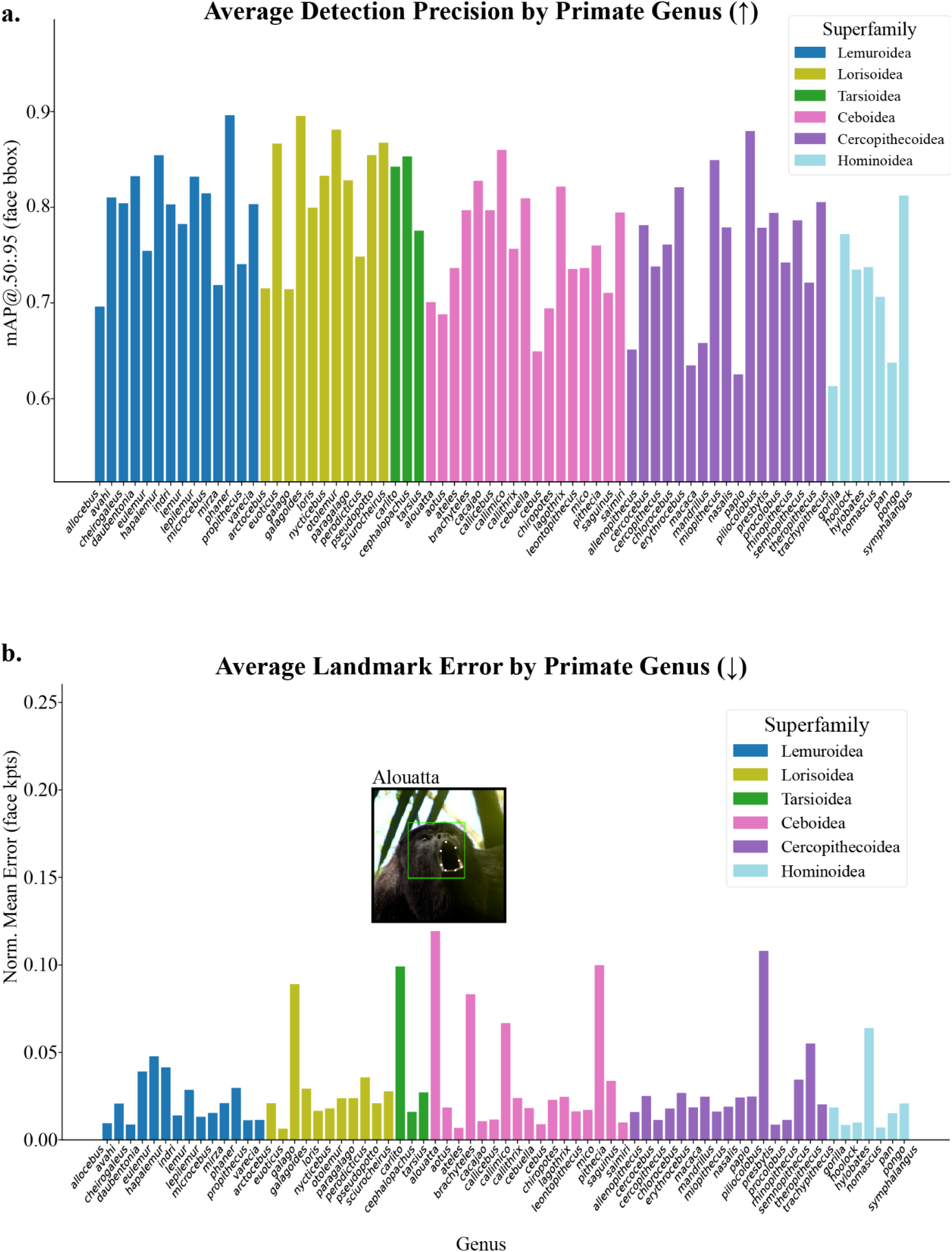
Face detection and facial landmark estimation performance by genus. Genus-level performance breakdown revealing heterogeneous model accuracy across the taxonomic diversity of PrimateFace. A. Mean average precision (mAP@0.50:0.95) per genus for face detection, showing variation that likely reflects differences in data curation, image quality, and ecological contexts across genera. A. Normalized mean error (NME) per genus for facial landmark estimation, demonstrating the challenges of achieving uniform performance across morphologically diverse primate faces. Performance variations highlight the importance of continued data collection efforts for underrepresented genera and provide guidance for researchers working with specific taxa about expected model accuracy.

**Extended Data Figure 3.**
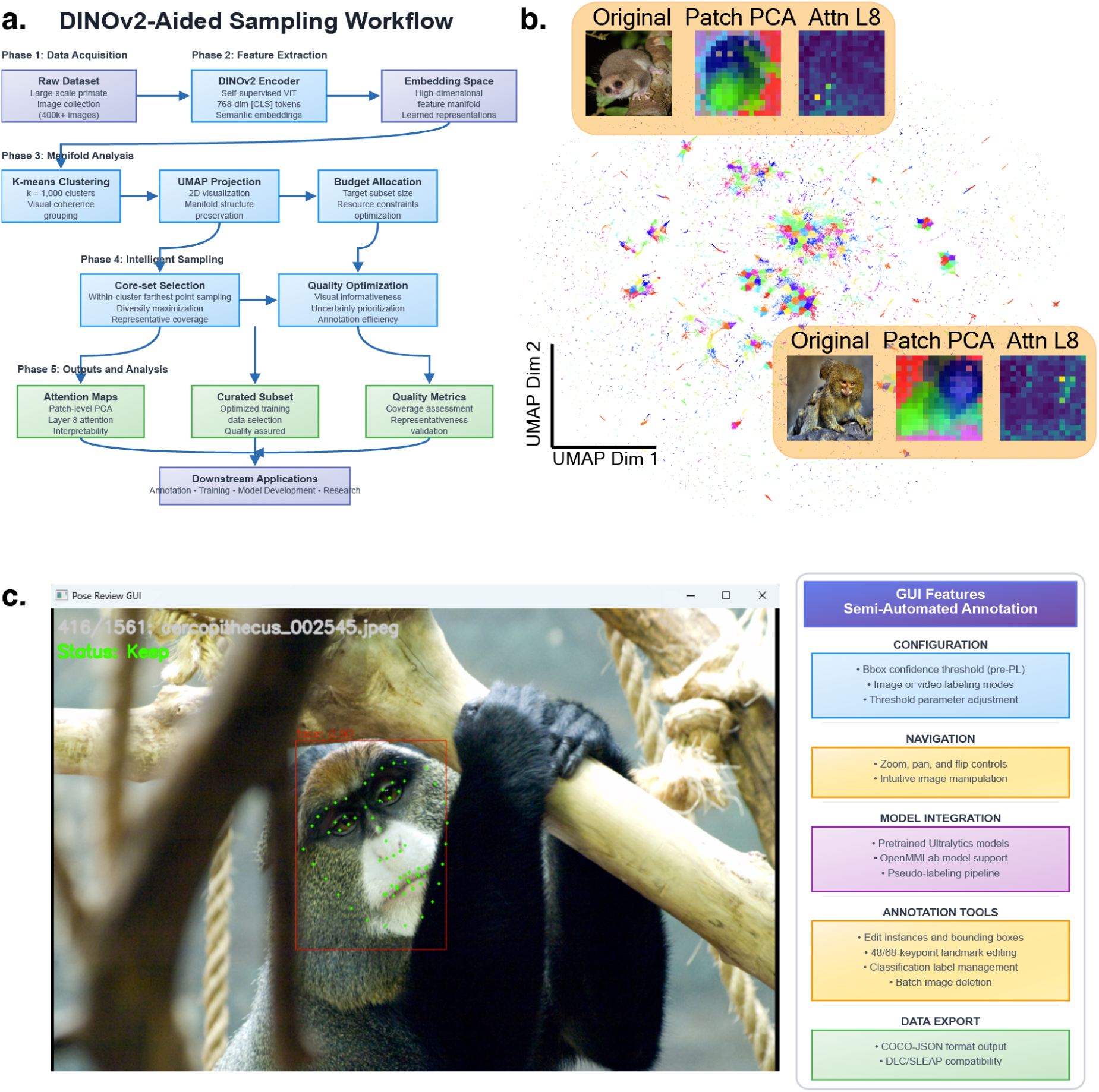
Scalable methods for dataset curation and annotation. A. Overview of DINOv2-aided efficient image sampling from larger datasets. A. UMAP projection of 400,275 primate face embeddings, extracted from the [CLS] token of a DINOv2-base model. Points are colored by K-means cluster assignments (k=1,000) to highlight visually coherent groupings. Insets: three example images with visualizations of the patch-level PCA (center) and Layer 8 attention (right), show semantic focus on facial regions. C. Custom GUI used for human-in-the-loop pseudo-labeling. Annotators iteratively refine bounding boxes and keypoints, guided by model predictions. This workflow enables efficient annotation of large-scale datasets by prioritizing visually informative and uncertain samples. Acronyms: DINOv2, self-supervised Vision Transformer; UMAP, Uniform Manifold Approximation and Projection; PCA, principal component analysis; CLS, class token; GUI, graphical user interface.

**Extended Data Figure 4.**
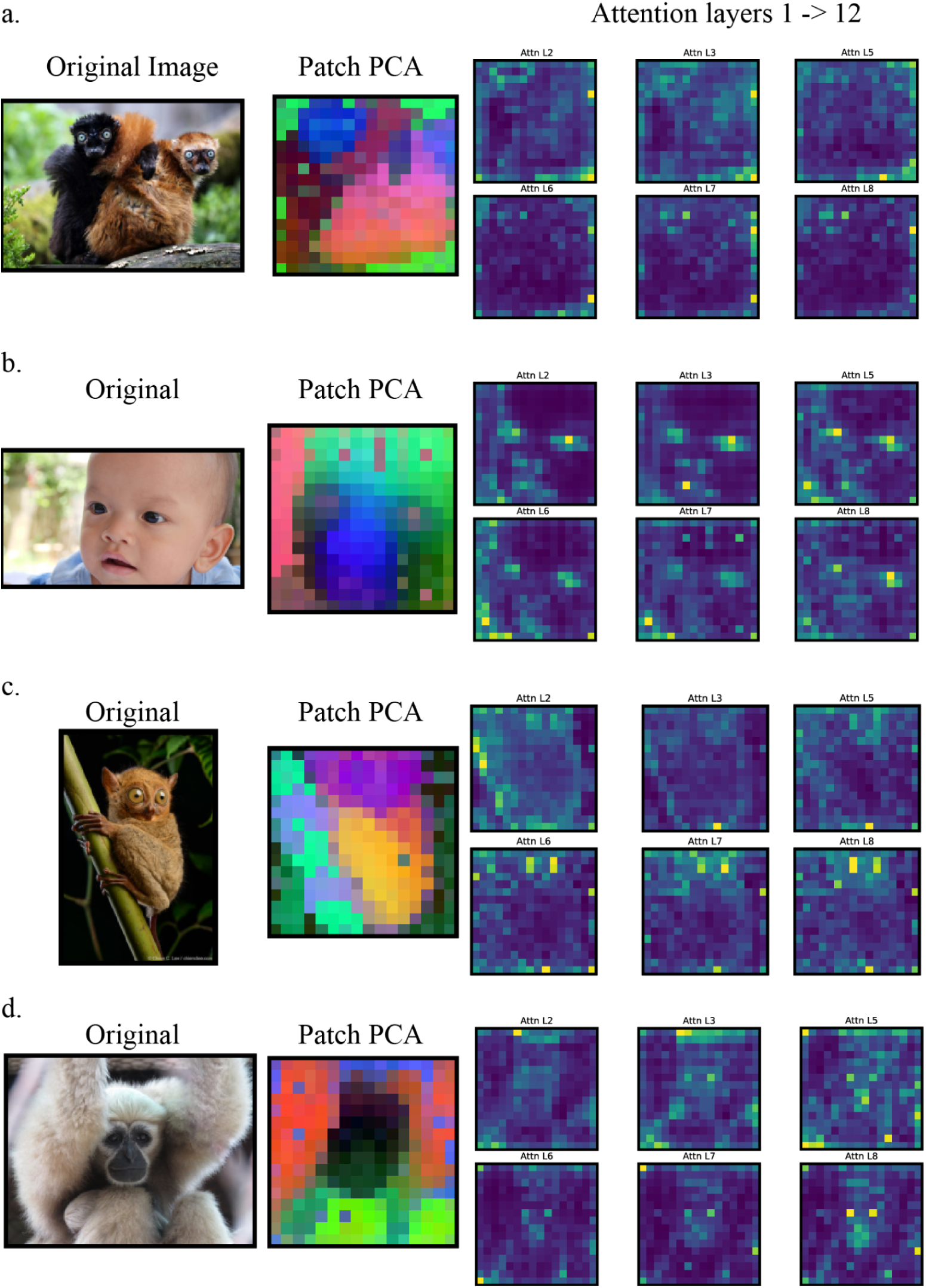
Attention heatmaps of DINOv2 features consistently localize salient facial regions. Each panel displays original image, PCA visualization of patch-level features, and attention heatmaps from 6 of 12 DINOv2-base attention layers. Heatmaps demonstrate that attention progressively focuses on semantically relevant areas, such as the eyes and snout, across layers. A. Blue-eyed black lemur (*Eulemur flavifrons*); A. Stock image of human infant (Adobe Stock); C. Horsfield’s tarsier (*Cephalopachus bancanus*); D. Hoolock gibbon (*Hoolock sp.*).

**Extended Data Figure 5.**
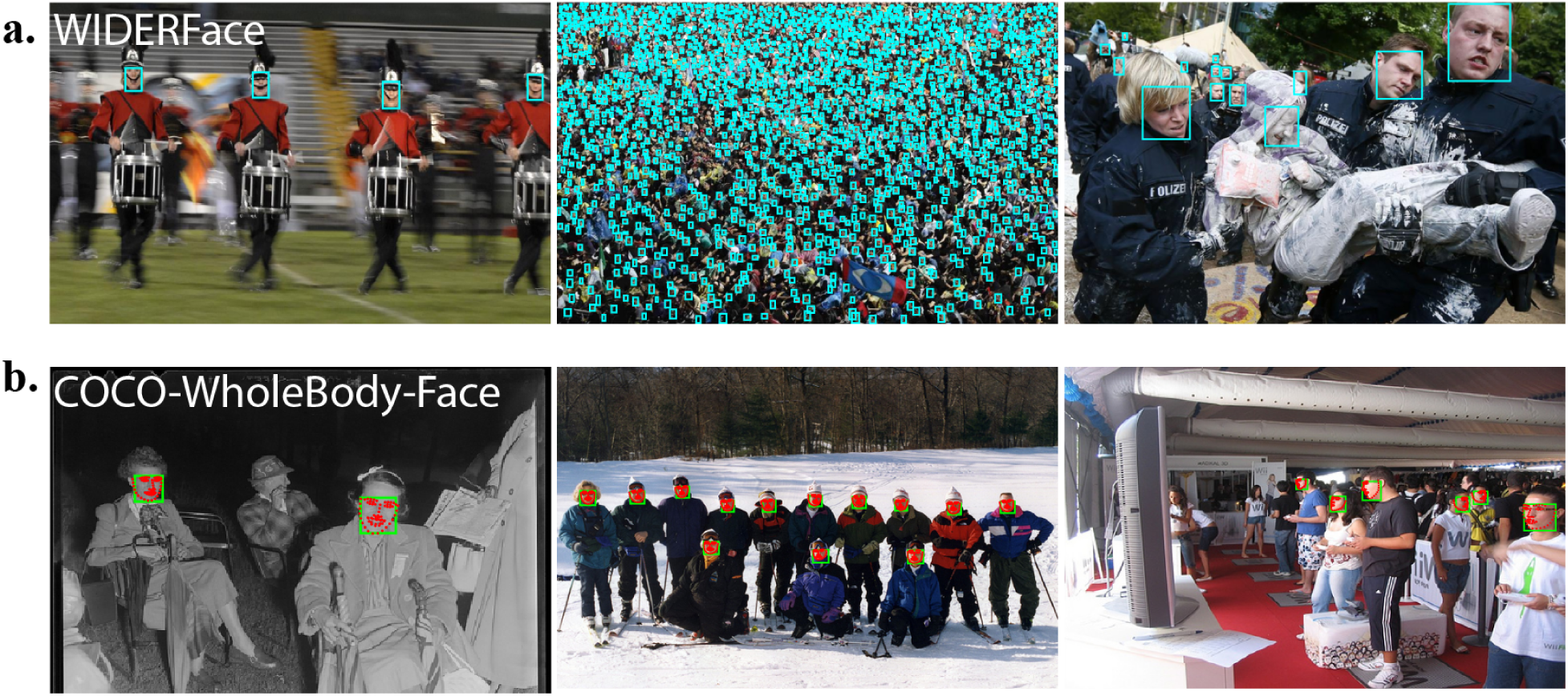
Human benchmark datasets for cross-domain evaluation. Representative images from standard human face analysis benchmarks used to assess PrimateFace model generalization. A. WIDERFace dataset examples showing challenging human face detection scenarios including blurry scenes, extremely crowded scenes, variable lighting, and scale variations that make this benchmark notoriously difficult for face detection models. Bounding boxes overlaid in light blue. A. COCO-WholeBody-Face dataset examples demonstrating diverse human facial poses and expressions used for facial landmark estimation evaluation. Bounding boxes overlaid in light green; facial landmarks in red. These benchmarks provide critical validation that PrimateFace-trained models maintain competitive performance on human faces despite being optimized for cross-species primate analysis, demonstrating the value of morphologically diverse training data for robust face analysis applications.

**Extended Data Figure 6.**
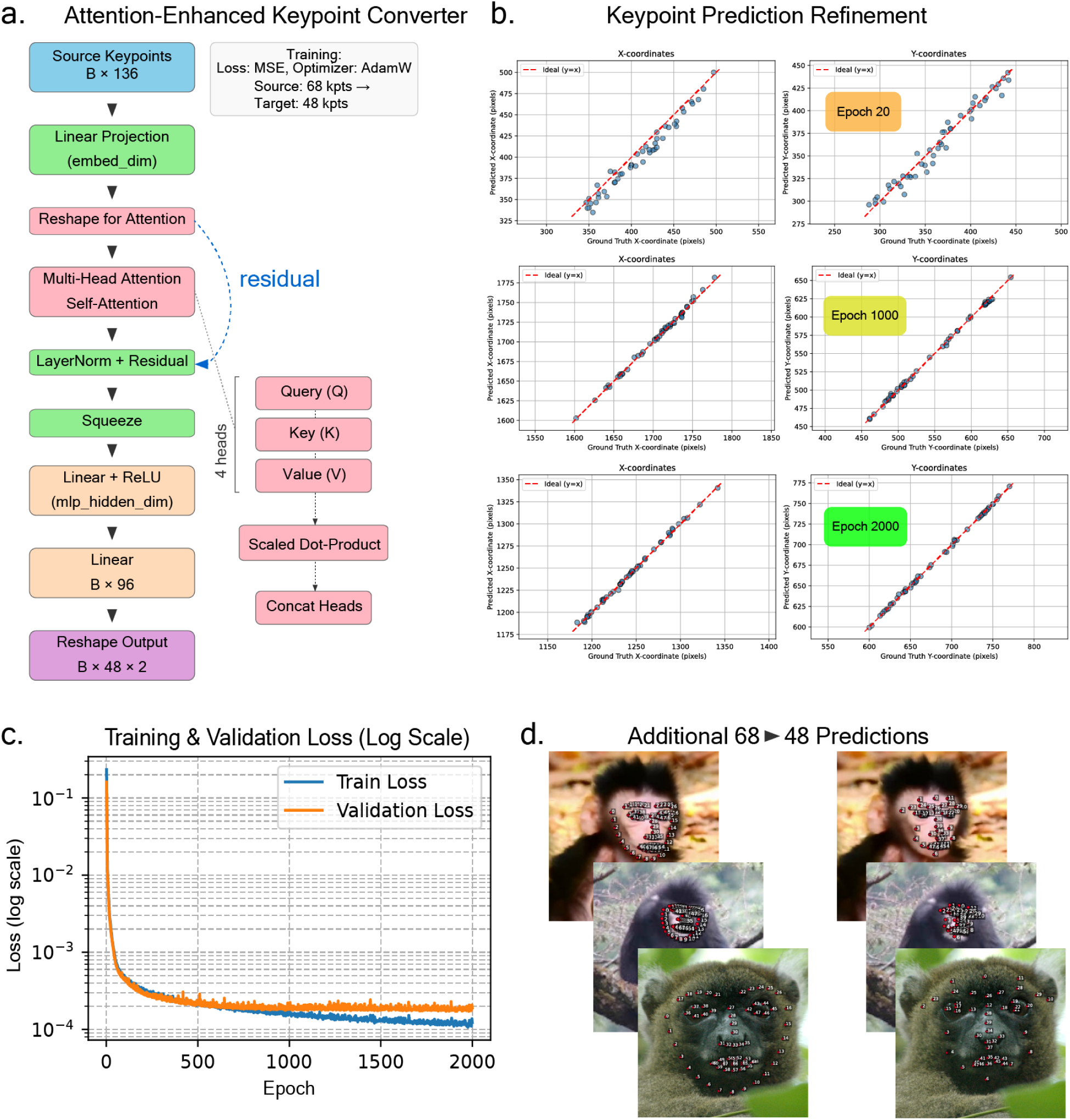
Landmark conversion model architecture and training validation. **A.** Architecture of the Attention-Enhanced MLP keypoint conversion model showing input 68-keypoint coordinates, attention mechanisms, and output 48-keypoint predictions. A. Scatter plots visualizing prediction error for x-coordinates (x-axis) and y-coordinates (y-axis) across training progression, with points representing individual landmark predictions and colors indicating training epochs (progression from top to bottom). C. Training and validation log-loss curves demonstrating model convergence over 2000 epochs. D. Additional examples of keypoint conversion predictions, showing input 68-keypoint configurations (human standard) transformed to output 48-keypoint format (PrimateFace custom) across diverse faces, demonstrating the model’s accurate translation between landmark schemes while preserving anatomical correspondence across species.

## Notes

https://primateface.studio/

### Competing Interest Statement

MLP is a scientific advisory board member, consultant, and/or co-founder of Blue Horizons International, NeuroFlow, Cogwear Technologies, Glassview, Lazul LLC, Almond Digital Health, elanah.ai, Mandala LLC, FeLiCiTi LLC, and NinetyPlay, and receives research funding from AIIR Consulting, Slalom Inc, Fox Media, Korn Ferry, Deloitte, and Glassview. All other authors declare no competing interests.

### Summary of Updates

Added additional author details e.g., co-senior author and correspondence email.

